# A cell type-resolved proteomic atlas of the human body

**DOI:** 10.64898/2026.05.26.727663

**Authors:** Caroline A. M. Weiss, Evelina Sjöstedt, Adhiraj Debnath, Shani Ben-Moshe, Lucas Diedrich, Martin Zwahlen, Marc Oeller, Maria Wahle, Emilio Skarwan, Tim Heymann, Denys Oliinyk, Elena Kratz, Andreas Metousis, Christian Braun, Heribert Schunkert, Fredric Johansson, Emma Lundberg, Kalle von Feilitzen, Cecilia Lindskog, Florian A. Rosenberger, Moritz von Scheidt, Mathias Uhlén, Matthias Mann

**Affiliations:** Department of Proteomics and Signal Transduction, Max Planck Institute of Biochemistry, Martinsried, Germany; Department of Protein Science, KTH Royal Institute of Technology, Stockholm, Sweden; Department of Neuroscience, Karolinska Institutet, 17177 Stockholm, Sweden; Institute of Legal Medicine, Faculty of Medicine, LMU Munich, Munich, Germany; Department of Cardiology, TUM University Hospital German Heart Center, Technical University Munich, Munich, Germany; Deutsches Zentrum für Herz- und Kreislaufforschung (DZHK), Partner Site Munich Heart Alliance, Munich, Germany; Department of Bioengineering, Stanford University, Stanford, CA, USA; Science for Life Laboratory, School of Engineering Sciences in Chemistry, Biotechnology and Health, KTH – Royal Institute of Technology, Stockholm, Sweden; Department of Pathology, Stanford University, Stanford, CA, USA; Department of Immunology, Genetics and Pathology, Uppsala University, Uppsala, Sweden; Department of Medical Biochemistry and Biophysics, Science for Life Laboratory, Karolinska Institutet, Solna, Sweden

**Author notes:** These authors contributed equally to this work (Florian A. Rosenberger, Moritz von Scheidt, Mathias Uhlén, Matthias Mann).

## Abstract

Proteins define what cells do, yet their systematic quantification across human cell types has remained out of reach. Using Deep Visual Proteomics on tissue from a healthy female donor, we built an atlas of 27 cell types across 14 tissues, quantifying two-thirds of all human protein-coding genes, with up to 8,500 per population. The proteome partitions bimodally into a universal core and highly specialized programs. Integration into the Human Protein Atlas Single Cell Resource enabled comparison of RNA and protein abundance at cell type resolution, revealing that concordance depends on pathway rather than cellular identity. This resolution uncovered cancer-testis antigens in oocytes invisible to bulk profiling. Our openly accessible resource provides a foundation for cell type-resolved proteomics in health and disease.

## Introduction

The human body comprises many distinct cell types that share essentially identical genomes yet each execute strikingly divergent functions essential for tissue homeostasis and organismal physiology (*1*, *2*). A contracting cardiomyocyte, a granulosa cell supporting an oocyte, and a metabolically active hepatocyte differ not in their DNA but in the proteins they deploy and disruptions to this finely balanced ensemble can result in severe disease. The proteome is therefore where cellular identity is actually written. A comprehensive catalogue of protein expression across human cell types would hence be foundational for understanding both normal physiology and pathological states.

Systematic efforts to map the molecular basis of human cellular identity have progressed through multiple scales. The Human Genome Project established the complete genetic blueprint (*3*, *4*) while subsequent transcriptomic atlases have since revealed the remarkable heterogeneity of gene expression across tissues and cell types. These range from bulk tissue resources such as GTEx (*5*) to cell type-resolved efforts including the Human Cell Atlas (*1*) and Tabula Sapiens (*6*). Yet transcript abundance is an imperfect proxy for the proteins that ultimately execute cellular function, and the magnitude and pattern of this divergence across human cell types has remained largely uncharted (*7*, *8*). Antibody-based imaging, pioneered by the Human Protein Atlas (HPA) (*9*, *10*), provides the most extensive spatial view of human protein expression data to date, with information for over 15,000 proteins across 45 tissues, but each protein must be addressed one antibody at a time.

Mass spectrometry (MS) offers a complementary capability: unbiased, high-throughput quantification of thousands of proteins simultaneously with excellent specificity. Early MS-based efforts mapped protein expression across human organs, yet bulk measurements average over the very cellular diversity that defines tissue function and the proteins most diagnostic of cell identity are often diluted below detection (*8*, *11*, *12*). Achieving cell type resolution by MS has remained technically challenging, requiring the isolation of defined cell populations from intact tissue while preserving sufficient material for proteomic analysis. As a result, fundamental questions about the protein-level organization of human cell biology remain open: which proteins are universal across cell types and which are truly cell-type-defining; how reliably does transcript abundance predict protein abundance once cell-type averaging is removed; and what biology lies hidden in rare or hard-to-isolate cellular populations? Recognizing this limitation, global initiatives such as the Human Proteome Project (HPP) (*13*) and the π-HuB consortium (*14*) have called for systematic mapping of the human proteome at cell-type resolution.

Here we address this critical gap, with a cell type-resolved human protein atlas mapping 27 distinct cell types across 14 tissues including cardiomyocytes and skeletal myofibers, neurons and astrocytes, hepatocytes, kidney tubular and glomerular cells, immune populations, and reproductive cells including oocytes and granulosa cells, all from a single healthy donor. This is enabled by Deep Visual Proteomics (DVP), a technology that combines AI-guided cell detection and classification with laser microdissection and ultra-high-sensitivity MS (*15*).

Integration with matched single-cell transcriptomes through the Human Protein Atlas Single Cell Resource then allows us to ask, for the first time at high resolution and scale, how the proteome is organized across human cell types and where it diverges from the transcriptome. Additionally, side-by-side comparison with matched bulk tissue proteomics data illustrates how biology with translational impact can sometimes only be revealed at single cell type resolution.

## Results

### A cell type-resolved proteomic atlas of human tissues

To build this atlas, we obtained formalin-fixed paraffin-embedded (FFPE) tissue sections from a healthy donor (24 years; BMI 23.4) at 15 hours postmortem, a window in which protein integrity remains well preserved (**Fig. S1**). Sections were processed through the DVP pipeline combining high-resolution immunofluorescence microscopy, AI-guided cell segmentation and laser microdissection, followed by MS-analysis on the latest generation ultra-high-sensitivity MS Orbitrap Astral Zoom platform (*16*) (**Fig. 1A**, Methods). The single female donor design eliminates inter-individual variability in genetic background and lifestyle. Additionally, it provided access to reproductive-age ovarian and fallopian tube tissue, enabling us to profile reproductive cell types that remain underrepresented in proteomic studies.

**Fig. 1.**
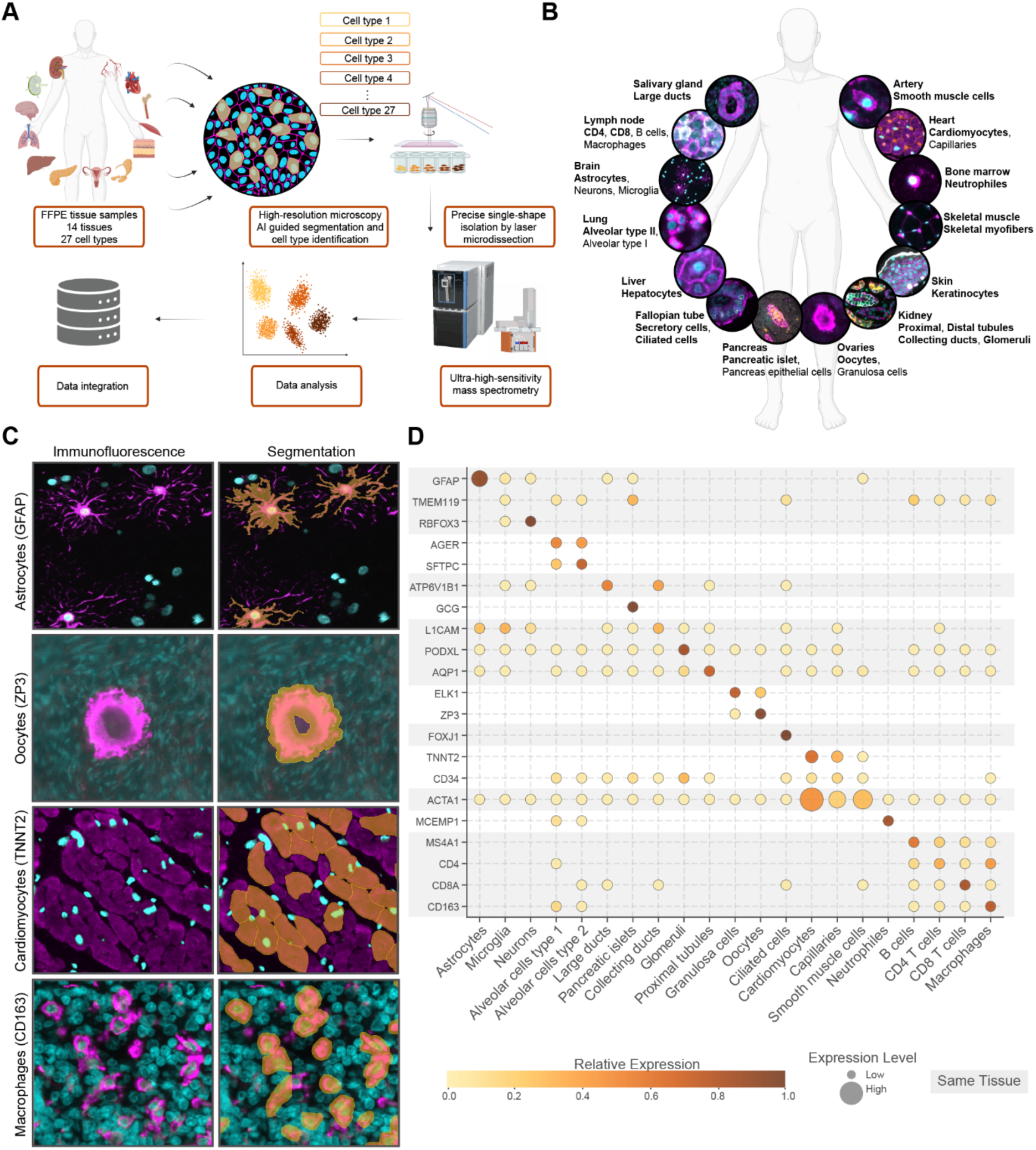
Deep visual proteomics enables cell-type-resolved proteomic mapping of human tissues. (**A**) Overview of the deep visual proteomics pipeline. Formalin-fixed paraffin-embedded tissue sections from 14 human tissues undergo high-resolution immunofluorescence microscopy, AI-guided cell segmentation and laser microdissection for precise single-cell isolation, followed by ultra-high-sensitivity mass spectrometry and data analysis plus integration with the Human Protein Atlas. (**B**) Illustration of the 27 characterized cell types profiled across 14 tissues. Representative immunofluorescence images are shown for selected cell types (bold) per tissue. (**C**) Representative immunofluorescence images (left) and corresponding segmentation masks (right) for astrocytes (GFAP), oocytes (ZP3), cardiomyocytes (TNNT2), and macrophages (CD163). Nuclei are shown in cyan, cell-type-specific markers in magenta, and segmentation contours in orange. (**D**) Dot plot showing expression of marker proteins used for cell identification across 20 cell types. Dot color indicates relative expression (proportion of total expression across cell types); dot size indicates absolute expression level; cell types obtained from the same tissue are indicated by alternating background color.

The 27 profiled cell types capture broad anatomical and functional diversity, spanning the central nervous system, cardiovascular, immune, metabolic, reproductive, epithelial, and musculoskeletal compartments (**Fig. 1B**). Within each tissue, the selected cell types represent essential specialized functions, from astrocytes maintaining neuronal homeostasis to cardiomyocytes driving cardiac contraction. For each, we developed customized segmentation pipelines guided by well-characterized marker proteins visualized through immunofluorescence (**Fig. S2**). For example, GFAP enabled identification of astrocytes, ZP3 delineated oocytes, TNNT2 marked cardiomyocytes and CD163 identified macrophages (**Fig. 1C**).

For most cell types, we collected five replicates of 100,000 µm² of cell type area by laser microdissection, with fewer replicates for rare populations. This corresponds to cell numbers below 100 for astrocytes, skeletal myofibers and oocytes replicates (complex architecture or exceptionally large) to about 1,000 for immune cells and some specialized epithelial cells (**Fig. S3A**). We found that the area-based approach best accommodated the morphological diversity in shape and size of cell types and enabled dataset-wide normalization using directLFQ (*17*), a label-free quantification method that aligns protein intensity distributions between samples, allowing quantitative comparison across cell types (**Fig. S3B**). The resulting MS-based proteomes confirmed the specificity of stained protein markers across cell types, validating that the DVP workflow resolves distinct cell type identities even within shared tissue contexts. (**Fig. 1D, S4**).

### Deep proteome coverage across protein classes and cell types

Our results cover 67% of all human protein-coding genes, containing nearly 14,000 protein groups, where proteins sharing peptide sequences are grouped for unique identification (**Methods**). The captured dynamic range, spanning 8.5 orders of magnitude, allowed us to detect highly abundant cell type-specific proteins such as MYH7 in cardiomyocytes and CPS1 in hepatocytes as well as ubiquitously expressed proteins like GAPDH and VIM, together with low-abundance regulatory factors such as the transcription factors GATA2 in collecting ducts and NR2E1 in neurons (**Fig. 2A**).To systematically assess our coverage, we examined detection rates across diverse protein classes listed by the Human Protein Atlas (**Fig. 2B, Fig. S5**). The core cellular machinery was covered almost completely, with 86% of all annotated enzymes and 87% of metabolic proteins captured, including all 30 components of the citric acid cycle. Likewise, we identified 90% of proteins deemed essential for cell survival based on genome-wide knockout screens (*18*). This coverage dropped to 61% for predicted membrane proteins, a class traditionally thought to be challenging for MS due to hydrophobicity and poor solubility.

**Fig. 2.**
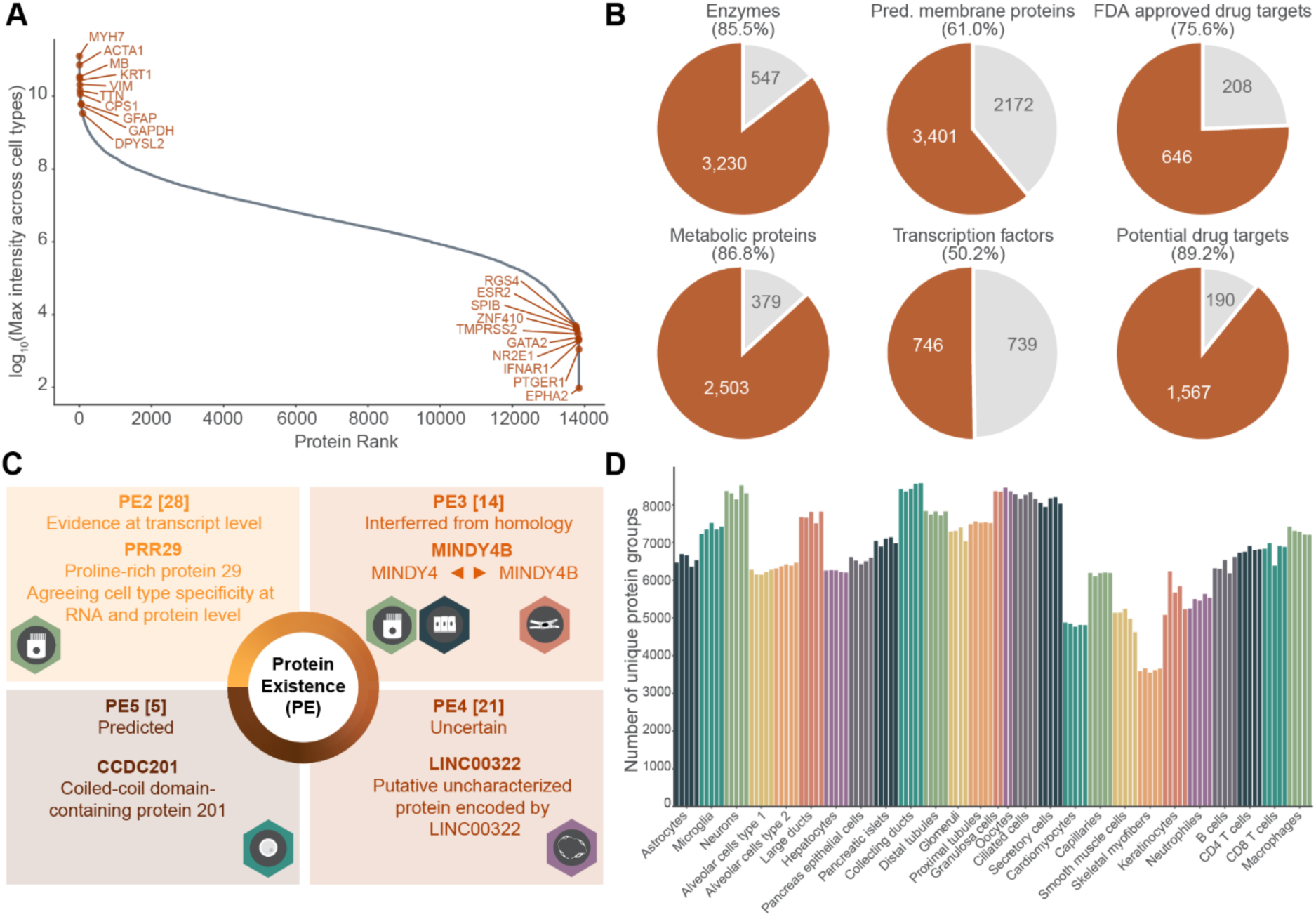
Deep proteome profiling reveals broad protein class coverage with cell type-specific depth. (**A**) Ranked protein intensity distribution across the complete dataset. Proteins are ordered by log₁₀ maximum intensity across all cell types. Selected high-abundance (left) and low-abundance (right) proteins are labeled. (**B**) Pie charts showing the proportion of detected (orange) versus undetected (grey) proteins for six protein classes annotated by the Human Protein Atlas (https://www.proteinatlas.org/humanproteome/proteinclasses). Percentages indicate detection rates. Numbers show absolute protein counts. (**C**) Distribution of 68 proteins detected by DVP that lack prior protein-level evidence in UniProt, categorized by protein existence level (PE2: transcript evidence; PE3: homology inference; PE4: predicted; PE5: uncertain). Representative examples are shown for each category; icons show their differential cell type distribution. Icons indicate cell type of detection. (**D**) Number of unique protein group identifications per cell type across 27 cell types. Individual replicates are shown.

Among membrane-associated subclasses, transporters and voltage-gated ion channels were captured at 76% and 62%, respectively. G protein-coupled receptors, whose seven-transmembrane topology poses a particular challenge for MS-based detection, were represented at 43.5% when excluding olfactory receptors. Notably, transcription factors, typically expressed at very low copy numbers and therefore among the most difficult targets for MS-based detection, reached 50% coverage. Finally, the dataset covers a substantial fraction of therapeutically relevant proteins, including 76% of FDA-approved drug targets (*19*) and 89% of potential drug targets (disease-associated proteins belonging to druggable protein classes not yet targeted by approved therapies) (*10*).

Only 6,670 protein coding genes remained undetected in the current dataset. Analysis using the GTEx database revealed that these predominantly map to tissues not represented in our study, with testis, pituitary and peripheral nerves the three top ones (**Fig. S6A-B**) (*20*). Overrepresented functions of this protein set primarily encompassed transcription factors, membrane proteins and sensory functions, the latter including olfactory receptors, which are not expected to be expressed in the tissues we sampled (**Fig. S6C-E**) (*21*). Together, these unsampled tissues and specialized protein classes were expected blind spots of our study design, rather than only due to dynamic range limitation.

Our data identified proteins for which, according to UniProt annotation, there was no prior experimental evidence at the protein level (**Suppl. Table S1**) (*22*). UniProt defines a protein existence (PE) evidence scale ranging from PE1 to PE5, with 91% of human protein-coding genes in PE1. Our data contained 28 proteins with transcript-level evidence only (PE2), 14 inferred from homology (PE3), 21 predicted proteins (PE4), and 5 proteins of uncertain existence (PE5; **Fig. 2C**). We manually validated the raw data for proteotypic peptides of these 68 proteins (**Fig. S7**, Methods). For instance, the proline-rich protein 29 (PRR29) was previously detected at transcript-level only with a high specificity to ciliated cells according to the Single Cell Resource of the HPA. DVP detected PRR29 at protein level and also recapitulate this cell type specificity in ciliated cells of the fallopian tube. In PE3 we found MINDY4B (PE3), predicted to be a deubiquitinase by homology. We detected and quantified it independently from its characterized homolog MINDY4, and also found a distinct expression specificity across cell types. Examples in PE4 and PE5 (uncertain and predicted proteins) included the long non-coding RNA-associated protein LINC00322 showing specific detection in capillaries, whereas the coiled-coil domain-containing protein CCDC201 was detected exclusively in oocytes (**Fig. 2C**). The cell type-resolved nature of our approach likely facilitated detection of such proteins by enriching for specific cellular populations rather than diluting them in bulk tissue lysates.

Oocytes yielded one of the deepest proteomes, consistent with maternal stockpiling of proteins for early embryonic development (*23*). In this cell type, we unexpectedly detected 35 proteins annotated as testis-specific by the HPA (*24*) (**Fig. S8B**). This may reflect shared germline expression programs that in previous work were masked by bulk tissue measurements due to the limited number of oocytes per sample and challenges in gaining access to ovarian samples from donors of reproductive age. Indeed, despite the testis-specific annotation there is previous experimental evidence for some of these proteins. For instance, the RNA-binding protein DAZL involved in translational regulation during gametogenesis was also reported at the protein level in human fetal oocytes and in adult mouse oocytes. However, direct protein-level evidence in adult human oocytes was lacking (*25*, *26*)

### The human proteome partitions into a universal core and cell type-specialized programs

Individual cell types differed by up to four orders of magnitude in their dynamic range of protein abundance, reflecting fundamental differences in cellular organization, protein content and the degree of cell type specialization (**Fig. S8C**). Cardiomyocytes exhibited the widest dynamic range, with highly abundant sarcomeric proteins dominating, consistent with the structural demands of cardiac contraction, and signaling and regulatory factors at trace levels. Within each cell type, we consistently identified known characteristic markers, further confirming our staining-driven cell type-selection approach. Beyond these expected cases, this resource identified additional candidate markers for specific cell types within their native tissue context, as exemplified by SMS as an oocyte-specific marker across tissues and SLC45A1 as a glomeruli-specific marker within the kidney, although also detected in other tissues (**Fig. S9)**.

The pronounced differences in proteome architecture that our data revealed raised the question of how proteins distribute across cell types and whether systematic functional relationships exist among them. To investigate this, we classified proteins by the number of cell types in which they were detected. Strikingly, the distribution followed a bimodal pattern, with proteins predominantly detected either in a single cell type or across all cell types, while intermediate detection frequencies were less common (**Fig. 3A**). Of the total of 1,766 proteins (12.8%) detected in only a single cell type, pathway enrichment analysis revealed characteristic functional specializations. This included cell cycle pathways in granulosa cells, lipid metabolism in keratinocytes, and drug and xenobiotic metabolism in hepatocytes, canonical functions of each cell type that confirm the biological specificity of the single-cell-type proteome, while providing a molecular inventory for them (**Fig. S10**).

**Fig. 3.**
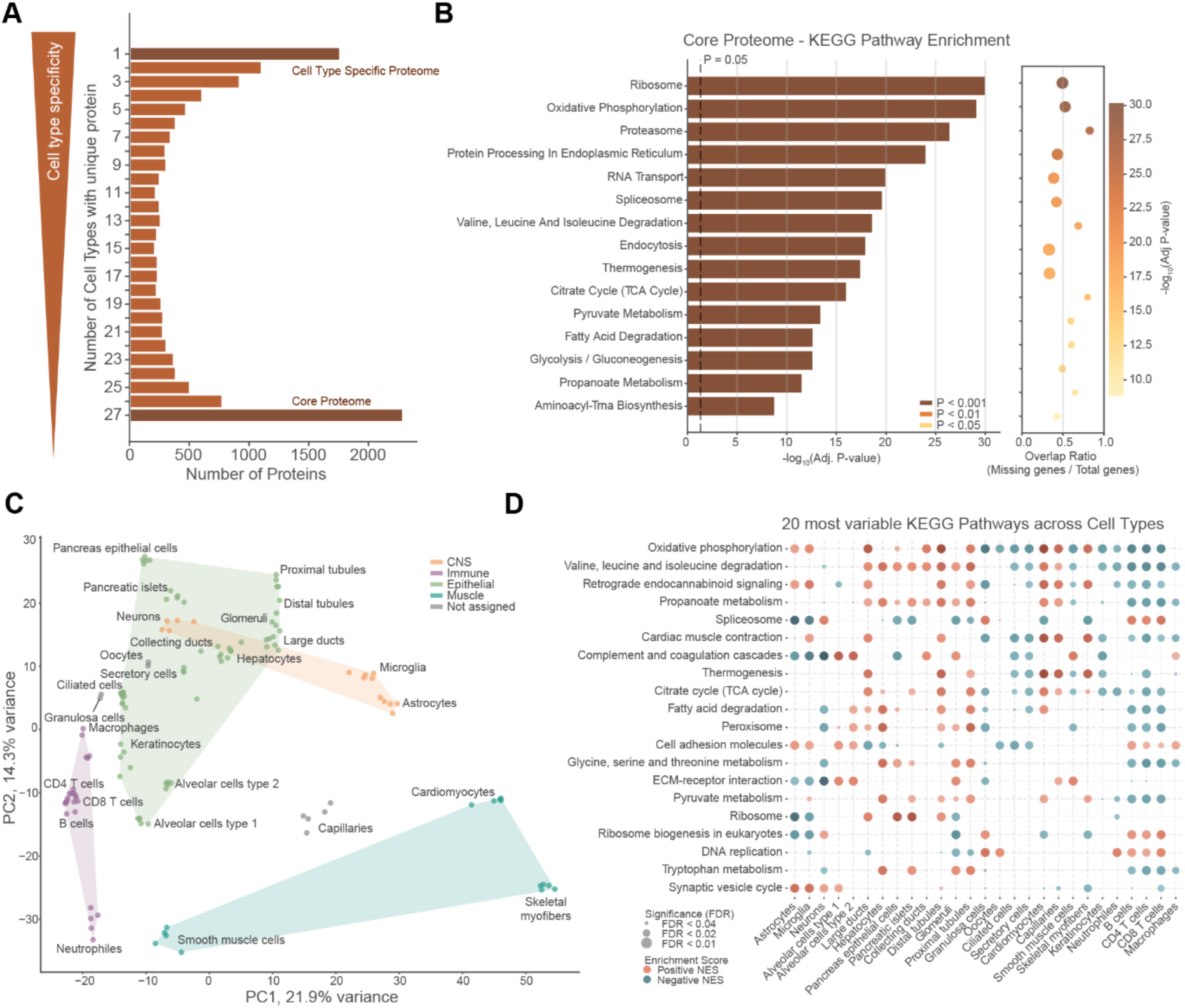
Proteome architecture reflects functional specialization across cell types. (**A**) Number of proteins detected in varying numbers of cell types (y-axis). Cell type-specific proteins (detected in one cell type) and core proteome (detected in all 27 cell types) are indicated. (**B**) KEGG pathway enrichment analysis of the core proteome. Bar length indicates −log₁₀(adjusted P-value) and bar color denotes significance level. Dot plot (right) shows the overlap ratio of detected genes to total pathway genes. (**C**) Principal component analysis of the core proteome across all cell types. Functional category annotation is color-coded (CNS, muscle, immune, epithelial, not assigned). (**D**) KEGG pathway enrichment analysis across 27 cell types. The 20 pathways with greatest variance across cell types are displayed. Dot color indicates enrichment direction (orange: positive normalized enrichment score, teal: negative). Dot size indicates significance level (FDR).

At the other extreme, our data defines a core proteome of 2,287 proteins (16.6%) across all 27 cell types. This is enriched for fundamental functions required for cellular survival, including energy metabolism (oxidative phosphorylation, glycolysis), protein synthesis (ribosome) and proteostasis (proteasome, protein processing) (**Fig. 3B**). Thus, the human proteome partitions into two principal classes: a universally expressed core and highly specialized cell type-specific programs, with relatively few proteins occupying intermediate ground.

We next asked whether proteome similarity among cell types reflects shared tissue of origin or shared cellular function. Principal component analysis (PCA) of the core proteome showed that replicates of the same cell type clustered tightly together and that cell types grouped by functional category rather than solely by the tissue from which they were isolated (**Fig. 3C, Fig. S11A,B**). Muscle cells separated distinctly along PC1 and immune cells clustered together on the opposite end, each spanning different organs yet converging in proteome space. CNS cell types formed a distinct group, with neurons closer to the pancreatic islets likely due to shared neuroendocrine proteins. In contrast, epithelial populations occupied a broader region, consistent with the diversity of specialized functions these cells perform across tissues. Analysis of shared proteins further supported these observations, with the largest proteome intersections formed by functionally related cell types (**Fig. S11C**).

Gene set enrichment analysis (GSEA) performed for each cell type revealed expected functional differences between cell types. We focused on the 20 most variable pathways to investigate these specializations (**Fig. 3D**). DNA replication was enriched in proliferative populations including immune and reproductive cells, synaptic vesicle cycle in CNS cell types and oxidative phosphorylation in cardiomyocytes and skeletal myofibers. Cell adhesion molecules showed enrichment in specific epithelial and immune populations. To quantify this functional commitment, we calculated the proportion of each proteome assigned to these 20 diverse pathways (**Fig. S12**). This fraction varied strikingly, from approximately 10% in keratinocytes to over 75% in cardiomyocytes and kidney tubular cells. This illustrates that some cell types dedicate the vast majority of their proteome to a small number of specialized functions while others maintain a more balanced allocation. These proteome allocation patterns mirrored the functional groupings observed by PCA at a quantitative level, reinforcing that shared function, rather than shared anatomy, is the primary organizer of cell type identity at the protein level.

### RNA-protein divergence follows pathway-level rules across cell types

We reasoned that our cell type-resolved proteomic atlas would enable the systematic assessment of the agreement of RNA with their corresponding proteins across matching cell populations, a comparison that has previously been limited to bulk tissues or cell lines or lacked the scale of a pan-organ analysis (*8*, *27–29*). To this end, we integrated our MS-based proteomes with single-cell and single-nuclei RNA sequencing (scRNAseq and snRNAseq) data from independent, unmatched donors provided by the Human Protein Atlas (**Methods**). For each of the 27 cell types, we correlated the DVP proteome with all RNA clusters from the corresponding tissue and selected best-matching clusters as the transcriptomic counterpart (**Fig. 4A, Fig. S13, Suppl. Table S2**). While some cell types yielded straightforward one-to-one matches, such as oocytes and granulosa cells in ovary, most required integration of multiple RNA clusters, as exemplified by hepatocytes and the immune cell populations.

**Fig. 4.**
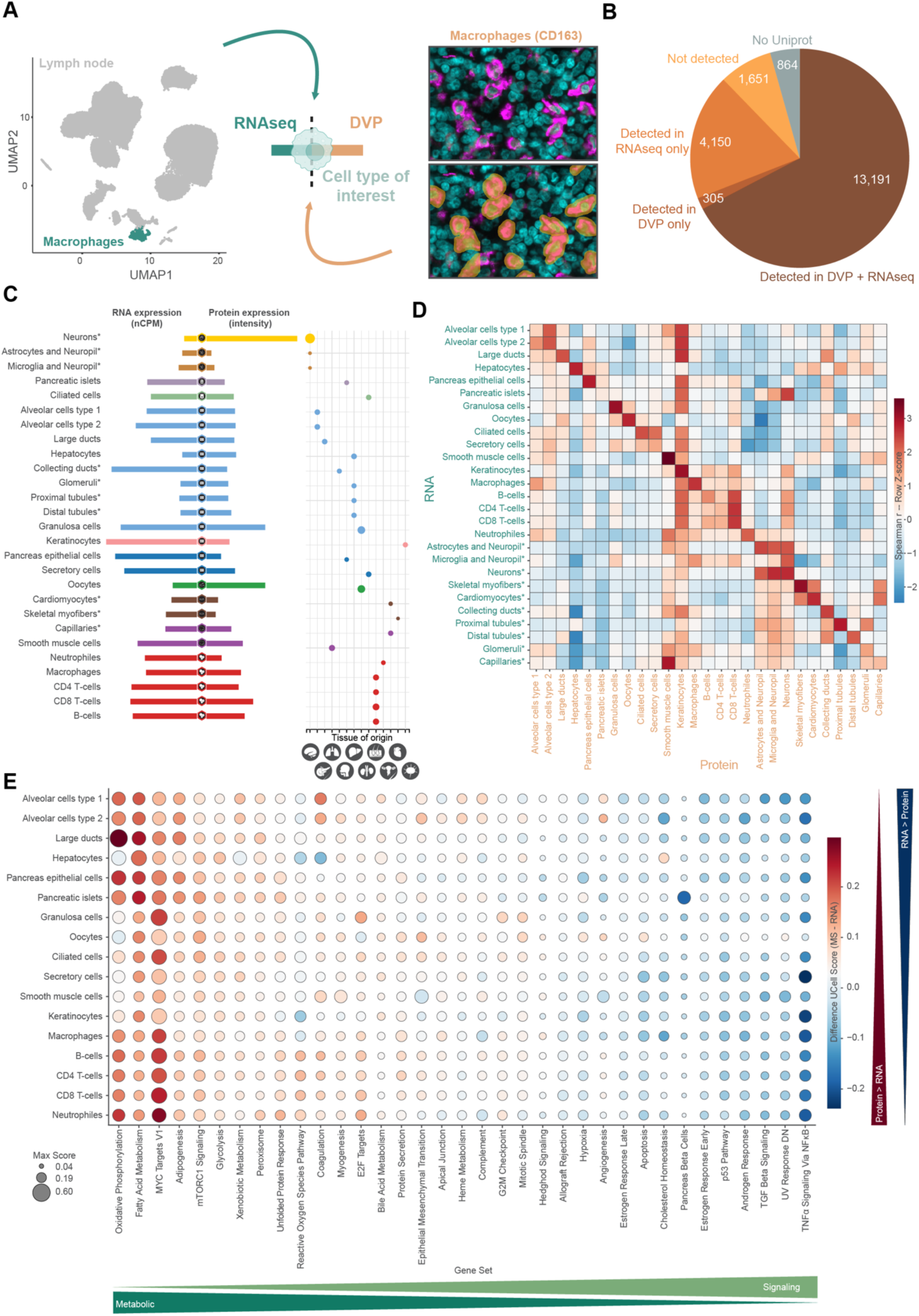
Cross-modality mapping uncovers cell-type specificity patterns across transcriptomic and proteomic layers. (**A**) Schematic of data integration strategy. For each cell type analyzed by DVP, matching single-cell or single-nuclei RNA sequencing clusters were selected from the Human Protein Atlas. Lymph node macrophages are shown as example. (**B**) Distribution of all human protein-coding genes (Ensembl v.109) by detection status across modalities. (**C**) Representative visualization of one matched RNA and protein expression data example as integrated into the Human Protein Atlas (the transcription factor NONO is used as an example). RNA expression is shown in normalized counts per million (nCPM); protein abundance is shown as intensity. (**D**) Spearman correlation matrix comparing RNA and protein expression across all cell types. Heatmap color represents row-normalized z-scores. Asterisks denote cell types with single-nuclei RNA sequencing data. (**E**) Comparison of Hallmark pathway representation between RNA and protein datasets using UCell scores. Pathways were filtered to include only those with a maximum UCell score exceeding 0.2 in at least one cell type. Dot color indicates the difference between protein and RNA UCell scores (red: higher at protein level; blue: higher at RNA level). Dot size indicates the maximum UCell score across both modalities. Pathways are ordered by mean difference across cell types. Higher representation of metabolic vs signaling pathways is indicated by green arrows. Cell types analyzed by snRNAseq are indicated by an asterisk.

Of the 20,162 protein coding genes in Ensembl v109 (current version used by the HPA v25), 864 lack matched UniProt identity, thus could not be annotated as detected by MS. For the 19,300 remaining ones, we found 13,191 in both RNA and MS datasets, 4,150 exclusively by RNAseq, 305 exclusively by MS, and 1,651 in neither modality (**Fig. 4B).** We have integrated this matched dataset into the Human Protein Atlas Single Cell Resource, providing an openly accessible platform for cell type-resolved RNA-protein comparison across human tissues (**Fig. 4C**).

We next asked how well transcript levels predict protein abundance at cell type resolution. Two caveats apply: the RNA and protein measurements derive from different biological samples, and the two platforms differ in sensitivity which attenuates observed correlations. Spearman correlations between matched transcriptome-proteome pairs ranged from 0.05 in capillaries to 0.37 in hepatocytes and smooth muscle cells (**Fig. S14A**). Crucially, comparison of each transcriptome against all 27 proteomes revealed that RNA clusters most frequently showed the highest correlation with their corresponding proteome, confirming the biological coherence of the pairings (**Fig. 4D**). scRNAseq datasets had much higher correlations than single-nuclei datasets in this matching (**Fig. S14B,C)**, reflecting the reduced dynamic range of nuclear-only RNA capture and the increased likelihood of cytosolic RNA to be translated.

To systematically compare RNA-protein abundance differences at the pathway level, we applied UCell scoring, a rank-based pathway enrichment method, to both datasets using all Hallmark pathways (*30*). Higher UCell score indicates higher relative abundance of the pathway members in the sample. The scores were calculated for each cell type on proteome and transcriptome level independently. Strikingly, individual pathways showed consistent RNA-protein differences across all cell types when calculating the difference between protein and RNA UCell scores for each pathway in each cell type (scRNAseq only; **Fig. 4E**). When we ordered pathways by their average difference, metabolic pathways, including oxidative phosphorylation, fatty acid metabolism and glycolysis, showed consistently higher representation at the protein level, while signaling pathways, such as TNFα signaling via NFκB and p53 pathway, were more strongly represented at the transcriptome level. Taken together, our data reveal predictable, pathway-level rules governing when transcript abundance does and does not reflect protein levels, pointing to distinct regulatory layers that predominate for different functional classes shared across tissues. The pathway ordering in **Fig. 4E** therefore serves as a practical guide to modality concordance and a starting point to dissect where regulation occurs along these axes.

### Transcriptomic and proteomic cell type specificity diverge in a functionally structured and translationally relevant manner

Because cell type identity is defined by selective gene and protein signatures, RNA–protein concordance at the level of individual genes becomes a central question. We find that this relationship is not uniform but varies systematically depending on the functional context of the gene. To investigate this, we merged cell types with very similar profiles, alveolar cell types 1 and 2 into alveolar cell types, ciliated and secretory cells into fallopian tube epithelial cells, and CD4 and CD8 T cells into T cells, reducing the set to 24 cell types.

Because the nature and magnitude of discordance varied widely, we employed three complementary metrics to capture different aspects of RNA-protein concordance (**Methods**). Beyond the established HPA categorization, we introduced a detection agreement score (A_Det_) that quantifies in how many cell types RNAseq and MS concur on whether a gene/protein is detected. Furthermore, the expression agreement score (A_Exp_) captures whether relative abundance is above or below a gene’s own median across cell types between both modalities. Both scores are necessary because pure detection as well as expression levels must be considered to fully interpret concordance. For instance, a protein may be identified across all cell types in both modalities yet show entirely divergent relative abundance patterns. GAPDH, classified as low specificity by HPA, illustrates how A_Det_ and A_Exp_ complement each other (**Fig. 5A**). It is detected across all 24 cell types (A_Det_ = 1.0, indicated by the matching triangles), however, A_Exp_ reveals that expression patterns differ substantially between RNA and protein (A_Exp_ = 0.33). Across all matched pairs the two scores have similar means (A_Det_ = 0.64, A_Exp_ = 0.65) but show markedly different distributions, indicating that characterizing RNA-protein relationships requires both detection and relative expression (**Fig. S15A**).

**Fig. 5.**
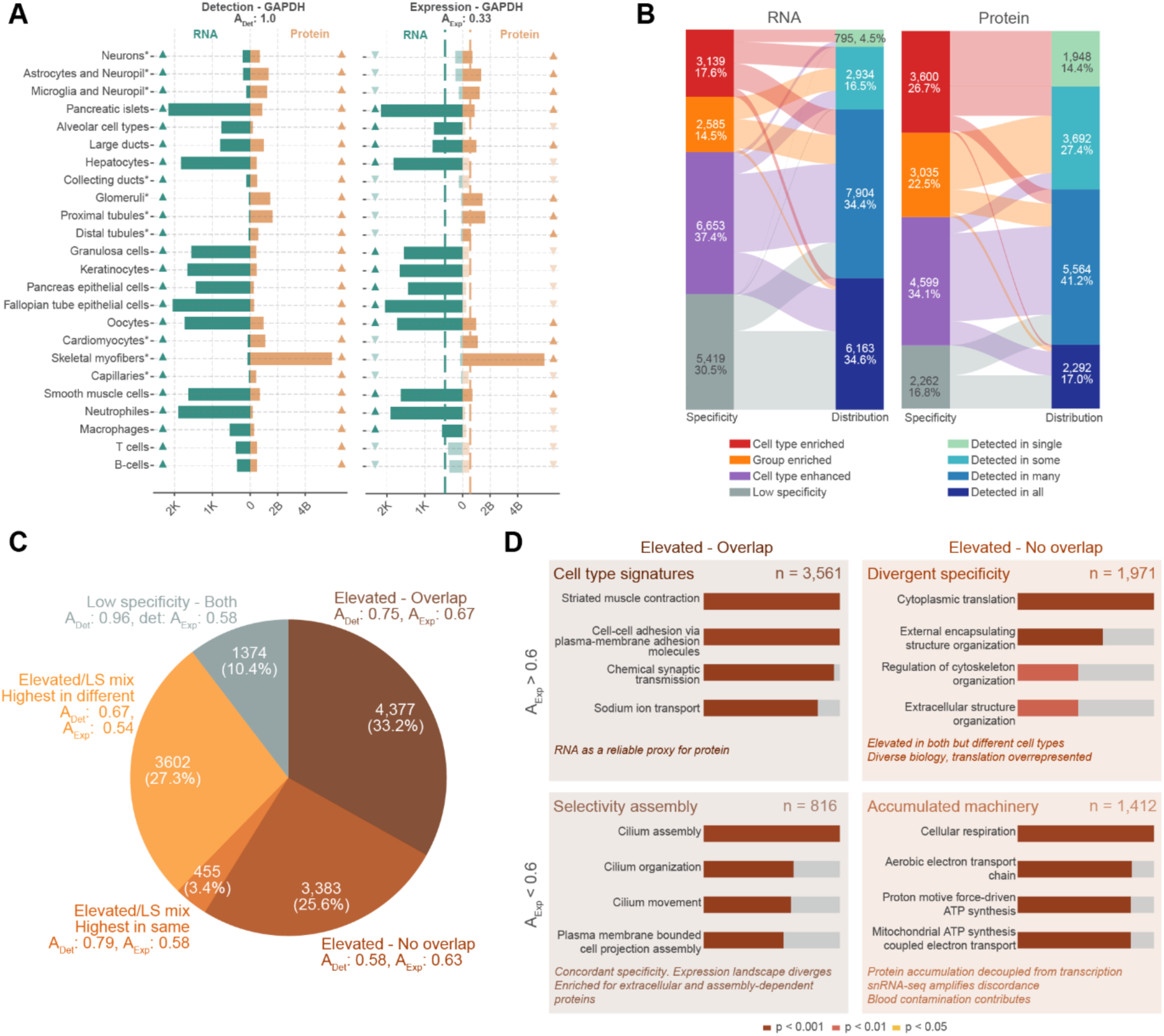
Concordance and divergence of cell type specificity between transcriptome and proteome. **(A)** Agreement scores quantifying RNA-protein concordance per gene across cell types. The detection agreement score (A_Det_) measures the proportion of cell types in which RNA and protein agree on whether a gene is detected. The expression agreement score (A_Exp_) measures the proportion of cell types in which both modalities agree on relative expression level (above or below the per-gene median). Both scores range from 0 to 1. GAPDH is shown as example (A_Det_ = 1.0; A_Exp_ = 0.33). RNA expression is shown in normalized counts per million (nCPM); protein abundance is shown as intensity (K = thousands; B = billions). Cell types analyzed by snRNAseq are indicated by an asterisk. (**B**) Transitions between category (left) and distribution (right) for RNA and protein datasets, classified according to Human Protein Atlas criteria. (**C**) Overlap of elevated status between RNA and protein for matched gene-protein pairs. Elevated overlap: elevated in the same (or some shared) cell types in both modalities; no overlap: elevated in different cell types. Elevated/LS mix indicates cases where one modality shows elevated and the other low specificity, with highest expression either in the same or different cell types. Median phi values within the group are labelled. **(D)** Overrepresentation analysis of genes stratified by elevated overlap status and expression agreement score. Genes with elevated status in both modalities are divided into those elevated in shared cell types (left, elevated overlap) and in different cell types (right, elevated no overlap), and further split by high (A_Exp_ > 0.6, top) and low (A_Exp_ < 0.6, bottom) expression agreement. Top four enriched biological process terms are shown for each group. Bar color indicates significance level.

Our HPA categorization describes both specificity and distribution patterns, applied here to both datasets (**Methods, Suppl. Table S3)**. Three proteins illustrate the spectrum across specificity categories. **(Fig. S15B**); GZMK, a cell type enriched serine protease restricted to T cells in both modalities; AGR3, group enriched across ciliated and alveolar cell types and TJP1, a cell type enhanced tight junction protein with broader expression profile that includes highest detection in renal glomeruli in both modalities. Globally, MS classified more proteins as elevated (any of the three specificity levels) than RNAseq, and distribution (categories that describe the detection frequency) classes shifted correspondingly, with RNAseq assigning more proteins to broader detection categories (**Fig. 5B**). Cross-tabulation of distribution classes confirmed this asymmetry (**Fig. S16A**): proteins shifted toward tighter distribution in MS rather than falling along the diagonal consistent with both lower per-cell-type sensitivity of MS and higher transcriptional noise in RNA inflating apparent detection spread. Note that, despite previously mentioned poor overall correlation on cell type level, very few individual cases show strong mismatch, with only 10 genes/proteins detected in all cell types according to MS while detected in single by RNAseq. Of the elevated proteins, 33% had overlapping specificity between RNA and protein, while 26% were elevated in both but in non-overlapping cell types. A further 31% switched between elevated and low-specificity categories across modalities (**Fig. 5C and Fig. S16B**). This incomplete concordance has translational implications, as a notable fraction of FDA-approved and potential drug targets are assigned different cell type specificity depending on the modality (**Fig. S16C**).

Combining specificity overlap with expression agreement A_Exp_ enabled further stratification into four classes (**Fig. 5D and Fig. S16D**). The largest comprised elevated proteins with both concordant specificity and high expression agreement (n = 3,579), enriched for identity-defining programs such as striated muscle contraction, cell-cell adhesion and chemical synaptic transmission. For these programs, RNA reliably predicts protein levels, as illustrated by MYOZ3 showing specific expression in skeletal myofibers in both modalities. At the opposite extreme, proteins with neither concordant specificity nor expression (n = 1,419) were enriched in mitochondrial respiration and translation, a pattern that persisted when restricting analysis to scRNAseq data. These long-lived metabolic proteins accumulate to steady-state levels that fundamentally decouple from transcriptional activity, illustrated by the nuclear-encoded component of complex I of the respiratory chain NDUFA1. Together, these data stratify RNA-protein pairs into distinct concordance classes with identity-defining programs showing strong agreement across modalities. However, a substantial fraction of elevated proteins, including FDA-approved and candidate drug targets, are assigned to different cell types depending on which modality is used, highlighting the necessity of a matched RNA-protein atlas for accurate cell type-resolved target identification and clues related to translatability across methods.

### A multimodal protein expression resource across human tissues

Bulk tissue proteomics complements the cell type-resolved atlas by providing broader organ context and directly exposing what is gained or lost without cellular resolution. We applied bulk MS profiling across all DVP tissues and six additional organs, including spleen and layers of the gastrointestinal tract (**Fig. S17A**). The resulting data are integrated into the HPA Tissue resource (**Fig. 6A**), adding an MS-based layer to the existing IHC and bulk RNA information enabling examination of any protein across all three modalities at bulk level.

**Fig. 6.**
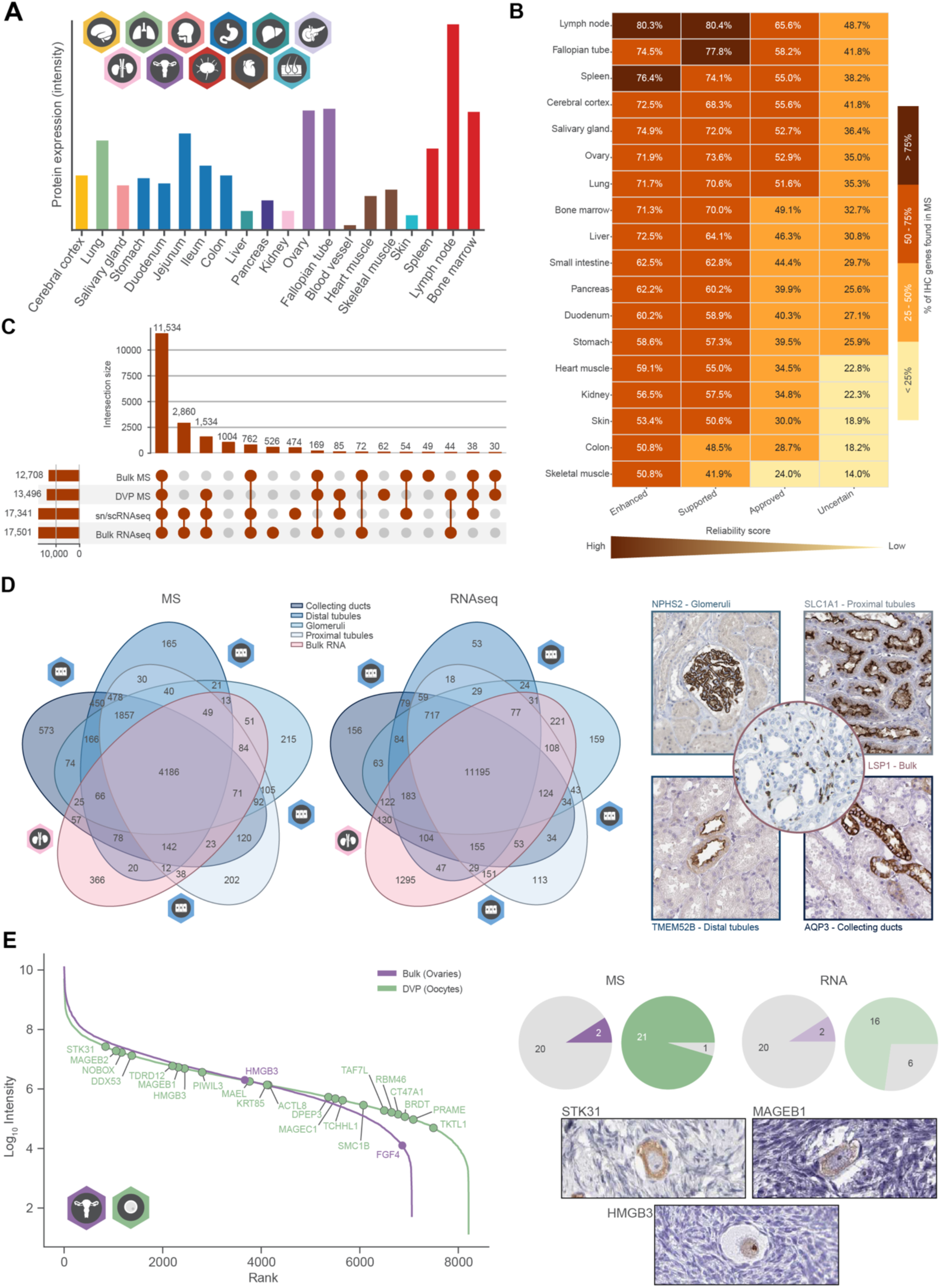
Bulk proteomics broadens atlas coverage and reveals protein signatures accessible only at cell type resolution. **(A)** Representative visualization of bulk MS data as integrated into the Tissue Resource of the Human Protein Atlas (the transcription factor NONO is used as an example). Protein abundance is shown as intensity. **(B)** Heatmap showing the percentage of HPA IHC-detected proteins recovered by bulk MS, stratified by tissue (rows) and HPA reliability score (columns; Enhanced, Supported, Approved, Uncertain). Color indicates detection rate. **(C)** UpSet plot showing the intersection of protein and transcript detection across four molecular profiling layers: bulk MS, DVP, sc/snRNAseq, and bulk RNA. Numbers indicate genes per intersection. **(D)** Venn diagrams comparing protein and transcript detection across four kidney-related datasets by MS (left) and RNAseq (right). Selected proteins exclusively detected in a single cell type/ bulk by MS are shown with representative images for NPHS2 (glomeruli), SLC1A1 (proximal tubules), TMEM52B (distal tubules), AQP3 (collecting ducts), and LSP1 (bulk). **(E)** Ranked protein intensity distributions for bulk ovarian tissue and DVP-profiled oocytes (left). Cancer-testis antigen (CTA) detection across bulk MS, DVP, bulk RNA, and single-cell RNA sequencing shown as pie charts for MS and RNA modalities (right, top). IHC validation of STK31, MAGEB1, and HMGB3 in oocytes (right, bottom).

In the HPA, antibodies are manually assigned a reliability score reflecting confidence in the reported protein expression pattern, organized as a four-tier hierarchy: Enhanced, Supported, Approved, and Uncertain. If MS-based detection provides an unbiased reference, recovery rates should track this hierarchy. Indeed, bulk MS recovers 86% of Enhanced-, 84% of Supported-, 78% of Approved-, and 67% of Uncertain-annotated proteins with concordant spatial IHC staining and MS-based detection illustrated for representative proteins from the Enhanced and Supported tiers (**Fig. S17B-C**). This trend was consistent across tissues, although absolute recovery rates varied from high agreement in immune-rich tissues such as lymph node to lower agreement in skeletal muscle (**Fig. 6B**). The one in three Uncertain-annotated proteins with IHC signal but undetected by MS is consistent with known antibody limitations such as cross-reactivity, dependency on dilution factors or mis-targeting. In this way our data can serve as a guide for re-evaluation of antibodies in HPA.

These four molecular profiling layers (bulk MS, DVP, bulk RNAseq, and sc/snRNAseq) collectively cover 94.6% of the mappable 19,300 protein-coding genes. The shared core of 11,614 genes detected across all four modalities represents the robustly quantifiable fraction of the proteome, while 1,034 remained undetected in any dataset. Consistent with the higher overall gene detection by RNAseq over MS noted above, 3,415 genes were captured exclusively by RNAseq-based approaches, while there were few proteins detected only by MS (**Fig. 6C**). RNA-only genes were enriched for cytokine-cytokine receptor interactions and neuroactive ligand-receptor signaling, consistent with the lower sensitivity of MS for membrane-bound receptors and low-abundance signaling molecules. Genes absent in all modalities mapped strongly to sensory functions, primarily olfactory transduction, which are not expected to be expressed in the tissues sampled here (**Fig. 6C**).

We examined protein and transcript detection across kidney as a case study for the complementarity of bulk and cell type-resolved measurements. As seen across the full atlas, RNAseq detected more genes overall (11,195 shared across cell types and bulk) while MS, despite lower coverage, was more discriminating at the cell type level (**Fig. 6D**). This pattern reversed in bulk tissue, where RNAseq uniquely detected more genes, likely because low-abundance proteins from rare populations are diluted beyond MS detection in unsorted lysates. Cell type-specific proteins detected by MS included known markers validated by IHC (**Fig. 6D**), and when mapped to kidney snRNA clusters they showed strong biological concordance, with DVP profiles aligning to their respective RNA clusters. Notably, proteins specific to bulk kidney MS showed the highest agreement with B cells and papillary tip epithelial cells, two populations absent from the DVP kidney dataset (**Fig. S17C**).

### Cell type resolution uncovers cancer-testis antigens in the female germline

The comparison between bulk and cell type-resolved measurements was most strikingly illustrated by oocytes, where cell type resolution not only increases proteome depth but uncovered a biologically important protein class invisible to bulk profiling. DVP profiled oocytes to a depth of 8,200 protein groups, even exceeding the 7,000 identified in bulk ovarian tissue. This gain in depth was accompanied by a wider dynamic range, explained by less signal dilution by the stromal tissue mass surrounding the rare oocytes in the intact organ (**Fig. 6E and Fig. S18A**). Interestingly, we identified 22 cancer-testis antigens (CTAs) by MS in oocytes (*31*, *32*). The expression of CTAs in healthy adult tissue is confined to immune-privileged sites, primarily testis and placenta, but they are frequently re-activated and overexpressed in malignant tumors(*33*). Their detection in oocytes is notable, as most have not been previously reported in healthy female germline cells, raising the possibility that oocytes share broader germline expression programs with testis than currently appreciated. Of the 22 CTAs, 21 were detected by DVP in oocytes and only two in bulk measurements of ovaries (**Fig. S18B**). Similarly, 16 CTA were detected by scRNAseq but only two by bulk RNA measurements. We further confirmed three CTAs by immunohistochemistry illustrating the spatially restricted expression in oocytes (**Fig. 6E**). This ten-fold gain in CTA detection (21 by DVP versus 2 by bulk MS) underscores that a cell type-resolved atlas captures a fundamentally different and richer view of the proteome than bulk profiling alone.

## Discussion

We present a cell type-resolved proteomic atlas of the human body, quantifying nearly 14,000 proteins across 27 cell types. As the atlas resolves individual cell populations rather than bulk tissues, it captures protein expression patterns that are invisible when cellular identities are averaged. The complete data set is integrated into the HPA Tissue and Single Cell Resource creating a freely accessible, multi-modal reference enabling community-wide exploration across MS, RNA and antibody-based modalities (v25.proteinatlas.org/humanproteome/single+cell/dvp).

This atlas was enabled by the combination of DVP with the latest generation of ultra-high-sensitivity mass spectrometry (*15*, *16*). DVP isolates defined cell populations directly from intact FFPE tissue sections, capturing populations inaccessible to dissociation-based approaches, including tightly connected epithelial cells and exceptionally large cells such as cardiomyocytes and oocytes. Nevertheless, certain biological constraints required adaptation such as podocytes, which were analyzed as part of glomerular units due to their integration with the glomerular basement membrane, and neurons which were restricted to cell bodies given their extensive projections. The upstream workflow, encompassing immunofluorescence optimization and segmentation pipeline development for all 27 cell types, required months of development, whereas MS acquisition of the entire dataset was completed in days. This asymmetry reflects the one-time investment in optimizing the DVP pipeline to a defined cell type. To enable others to bypass this step, all optimized markers, staining protocols and segmentation strategies are made openly available.

The resulting proteomes achieved broad coverage across biologically and clinically relevant protein classes reaching 86% of annotated enzymes, 90% of essential proteins and 76% of FDA-approved drug targets. Moreover, the identification of 50% of all transcription factors and 61% of predicted membrane proteins demonstrates that state-of-the-art MS provides access to the most challenging parts of the human proteome. This depth, reaching up to 8,500 proteins within a single cell type, goes beyond cataloguing known biology, providing a systematic resource for nominating and validating novel cell type markers. Notably, 68 proteins previously known only from transcripts or computational prediction were detected for the first time at the protein level. Strikingly, proteome organization across cell types followed a nearly bimodal distribution with proteins predominantly detected either in a single cell type or universally, with relatively few occupying intermediate ground. Hence, these patterns reveal that the human proteome partitions into a universal core sustaining cellular life and highly specialized programs that define cell type functionality.

Integration with matched sc and snRNAseq data from the HPA enabled the comparison of over 13,000 RNA-protein pairs. Overall, RNAseq detected a larger number of genes, whereas MS more frequently assigned proteins to elevated, cell type-enriched expression categories. This discrepancy reflects fundamental differences between the modalities. RNAseq captures low-level transcripts, including low abundant transcripts that are of limited reliability, and whose detection does not necessarily imply functional relevance or proportional protein output (*34–36*). In contrast, MS-based proteomics provides high specificity but remains less sensitive toward low abundance proteins (*37*, *38*). These modality-specific constraints introduce biases, further compounded by biological factors such as differences in dynamic range, cell state-dependent regulation (*39*), variation in RNA-to-protein copy number ratios (*29*), and spatial decoupling whereby secreted and circulating proteins are detected distal to their site of transcription (*8*). Differences in protein half-life present a particular challenge, as long-lived proteins can persist well beyond the presence of their transcripts while short-lived proteins may not accumulate to detectable levels despite active transcription (*40*).

A strong global correlation between RNA and protein abundance was generally not expected, as the limited concordance between transcript and protein levels has been widely documented over the past decades (*8*, *12*, *35*, *41–43*). Indeed, the observed correlations of 0.05-0.37 were modest, with the lowest concordance consistently observed when comparing protein data to snRNAseq data. This aligns with known technical and biological limitations of nuclear transcript capture, such as the reduced dynamic range (*44–46*). For several tissues, snRNAseq nevertheless represents the only viable option, as tight cellular junctions in kidney and complex cell morphology in brain render single-cell dissociation approaches technically unfeasible (*45*, *47*, *48*). Nonetheless, transcriptomes mostly showed highest correlation with its own matched proteomes across all 27 cell types, confirming that the pairings retain biological coherence despite the modest global correlations. Strikingly, the global comparison revealed consistent pathway-level patterns across cell types with divergence between RNA and protein not being cell type-specific but instead followed functional trends. Metabolic pathways were systematically enriched at the protein level, consistent with the long half-lives and stable expression of metabolic enzymes, whereas signaling pathways were more prominently represented at the transcript level as they are subject to rapid turnover or fall below the current detection sensitivity of MS (*38*, *40*). That the relationship between transcript and protein abundance is shaped by pathway membership rather than cell type identity implies that post-transcriptional regulation operates according to functional demand, an organizing principle conserved across human cell types. In general, deciphering at which regulatory layer each protein is primarily controlled remains a central goal of molecular biology and a prerequisite for predicting protein abundance from transcript measurements. This atlas lays a foundation for this effort by indicating that RNA-protein relationships are shaped more by pathway membership than by cell type identity, suggesting regulatory logic that generalizes across cellular contexts.

The scope of this atlas enables moving beyond global trends, mapping transcript-protein concordance at the level of individual gene-protein pairs (*29*). Approximately one third of elevated proteins showed concordant cell type specificity between RNA and protein. However, a substantial fraction displayed non-overlapping cell type specificity, highlighting cases where

RNA is not a reliable proxy for protein abundance and instead reflects more complex regulatory relationships with potential reasons mentioned above. The highest concordance was observed for identity-defining gene programs, underscoring that transcriptomic data can reliably predict protein expression for cell type signatures. Together, these findings underscore the importance of enabling user-driven exploration of the dataset. The implementation of agreement scores and category overlap provides a framework for interpreting RNA-protein relationships, allowing users to assess when transcriptomic data can serve as a proxy for protein expression and to identify proteins subject to more complex regulation. This is particularly relevant given the increasing reliance on scRNAseq atlases to infer cellular function (*6*, *36*, *38*, *49–51*).

Bulk MS profiling across 20 tissues provided both additional organ coverage and a direct measure of what cell type resolution adds. DVP and bulk MS detected comparable total protein numbers, yet proteins expressed in rare or specialized cell populations were diluted beyond detection in bulk tissue lysates. The oocyte proteome illustrates this most powerfully. MS-based approaches identified 22 cancer-testis antigens in oocytes, a discovery largely missed in matched bulk tissue analysis. As proteins whose expression in healthy adults is otherwise confined to immune-privileged sites such as testis and placenta, and which are actively pursued as immunotherapy targets (*31*, *33*, *52*, *53*), their presence in the female germline adds important biological context to their reactivation in cancer and raises questions about the safety profiling of CTA-targeting therapies in female patients. Moreover, DVP identified in oocytes 35 proteins that, based on previous bulk RNA profiling, are currently annotated as testis-specific. The large majority of these proteins were absent from matched bulk ovarian measurements. Both findings highlight the historical underrepresentation of female reproductive tissue in proteomic studies and the need for its more systematic investigation. Beyond resolving cell type-specific biology, the bulk MS dataset constitutes an independent, unbiased measurement of protein expression across tissues, providing an orthogonal criterion to assess antibody reliability in the HPA (*54*). Indeed, MS recovery rates tracked the annotation hierarchy consistently across tissues. Importantly, the reliability-annotation of lower tiers (Approved and Uncertain) reflects the limited availability of comparative data for validating these publicly available IHC profiles, including guidance on suitable positive and negative controls, thus determining an optimal dilution factor. Therefore, this resource can serve as an orthogonal criterion to re-evaluate and re-optimize antibodies currently presented in the HPA.

This atlas derives from a single donor, which eliminates inter-individual variability but allows matched cross-tissue comparison, precluding assessment of population-level variation across sex, age, genetic background and disease state. These limitations are recognized and actively addressed by emerging consortia including the HPP (*13*) and π-HuB (*14*). Furthermore, the tissues sampled do not cover the full anatomical diversity of the human body and tissues absent from this study account for the majority of proteins not yet detected. Lastly, matched multi-omic profiling of the same samples using consistent technologies will be essential to fully resolve the relationship between transcript and protein abundance at cell type resolution (*55*).

This work establishes a quantitative, cell type-resolved map of the human proteome that sets a new standard for the depth and resolution at which protein expression can be systematically characterized across the human body. Three findings from this atlas illustrate how it extends beyond the resource itself. First, the human proteome partitions bimodally into a universal core sustaining cellular life and highly restricted programs that define cell type identity, with relatively little intermediate ground. Second, the relationship between transcript and protein abundance is organized by pathway membership rather than by cell type, a regulatory logic that generalizes across cellular contexts and provides a practical guide for when RNAseq can serve as a proxy for protein expression. Third, cell type resolution exposes biology that bulk profiling obscures, including cancer-testis antigens in the female germline with direct implications for the safety profiling of CTA-targeting immunotherapies. Together with matched transcriptomic data in the HPA, our measurements place 95% of human protein-coding genes. Therefore, we provide the scientific community with an openly accessible, multi-modal reference spanning MS, RNA and antibody-based evidence at both tissue and cell type level. The dataset and all associated protocols are designed to be expandable across donors, tissues and disease states, addressing biological and translational questions that bulk measurements cannot resolve. Cell type-resolved proteomics is no longer a technical aspiration but a deployable strategy, and this work provides the foundation to apply it systematically across the full diversity of human physiology and disease.

## Authors contributions

Conceptualization: CW, ESj, FR, MvS, MU, MM

Investigation: CW, AD, LD, MO, DO, EK, AM

Methodology: CW, T.H

Data curation: CW, ESj, SB-M, LD, MZ

Formal analysis: CW, ESj, SB-M, LD, MZ, MW, ESk, CL

Resources: CB, HS, MvS

Visualization: CW, ESj, LD, MZ, FJ, KvF, CL, MU

Supervision: EL, KvF, CL, MU, MM

Writing - original draft: CW, ESj, SB-M, FAR, MM

All authors read, revised and approved the paper.

## Acknowledgements

We thank our colleagues at the Department of Proteomics and Signal Transduction at the Max Planck Institute of Biochemistry, as well as our colleagues at the Center for Proteome Research in Copenhagen, for their input and support. In particular, we thank T. Nordmann, M. Thielert, C. Ammar, M. Zwiebel, M. Murgia, J. Mueller-Reif, L. Schweizer, M. Schwoerer, for valuable input. We are further grateful for technical assistance of Katharina Zettl. We further thank Atlas Antibodies AB, for providing access to antibodies.

We acknowledge the Cardiovascular Biobank of the TUM University Hospital German Heart Center, Technical University Munich, as well as the laboratory, and research personnel, especially Sabine Bauer, Ulrike Weiss and Sophia Trautbeck, involved in the conduct of the study. This study was supported by the Max Planck Society for Advancement of Science and the WCPR grant KAW2022.0318 from the Knut and Alice Wallenberg Foundation. Furthermore, MO was supported by the HORIZON-MSCA-2023-PF-01-01 project HETLEWY, no. 101151819. AM is supported by a PhD scholarship from the Onassis Foundation (Scholarship ID: F ZS 031-1/2022-2023) and by an interdisciplinary life science fellowship awarded by the Joachim Herz Stiftung. CL is supported by the Swedish Research Council (2022–02742). FAR is an EMBO postdoctoral fellow (ALTF 399-2021) and supported by departmental and career grants from Karolinska Institutet. MvS is supported by the CORONA Foundation (Junior Research Group Cardiovascular Diseases Grant S199/10085/2021), the Deutsches CHIP Register e.V. (www.chip-register.de), the German Centre for Cardiovascular Research (DZHK) and the DZG Innovation Fund (81X2200145, 81X2600520, 81X3600506, 81X2600534), the German Heart Foundation, the German Research Foundation (DFG, INST 95/1713-1 FUGG), Förderverein Deutsches Herzzentrum München e.V., and the MIRACLE Consortium funded by the European Union and European Innovation Council program (Horizon Grant agreement No. 101115381) and the EuroHeartPath Consortium funded by the European Union and the Innovative Health Initiative Joint Undertaking (Horizon Grant Agreement No. 101194785).

## Methods

### Donor characteristics and ethical approval

The tissues analyzed in this study were obtained from the MunIch cardiovaScular StudIes biObaNk (MISSION). The MISSION was established in 2019 as a large-scale human tissue resource comprising post-mortem specimens from more than 1,500 individuals aged 18–98 years with an approximately balanced sex distribution. The collection has a focus on cardiovascular disease and aging and includes, among others, whole blood, plasma, liver, myocardium, and coronary and carotid arteries. The tissues analyzed for the DVP atlas and bulk tissue extension of it were derived from a 24-year-old female donor (173 cm, 70 kg; BMI 23.4) without any known acute or chronic diseases and a post-mortem interval of 15 h. The cause of death was traumatic and unrelated to underlying pathology, resulting from an accident. Heart weight was 317 g and histological assessment revealed no detectable vascular sclerosis. Donor information was handled in a fully anonymized manner. All procedures involving human material were reviewed and approved by the Ethics Committee of the Technical University of Munich (approval ID 325/18S) and conducted in accordance with the Declaration of Helsinki and relevant national regulations.

### Tissue procurement and anatomical sampling

Post-mortem tissue collection was performed at the Institute of Forensic Medicine following a predefined anatomical sampling scheme developed for the female human proteomic body atlas. The sampled tissues represent central nervous system, cardiovascular, immune, metabolic, reproductive, epithelial, and musculoskeletal compartments relevant to atlas construction. Collected tissues included bone marrow, cerebrum, coronary artery, heart (myocardium), skeletal muscle, kidney, liver, lung, lymph node, pancreas, ovary, fallopian tube, salivary gland, and skin. Sample metadata, including anatomical location and post-mortem interval, were documented using standardized collection sheets. Tissue specimens designated for histological analysis were transferred into formalin immediately after excision.

### Fixation and paraffin embedding

All tissues analyzed in this study were processed as formalin-fixed paraffin-embedded (FFPE) samples. Specimens were fixed in 4% neutral-buffered formalin for 48 hours, followed by transfer into graded ethanol. Tissues were trimmed to approximately 5 mm thickness, dehydrated through graded ethanol, cleared in xylene substitute, and infiltrated with paraffin using an automated tissue processor (total dehydration and infiltration time approximately 17 h). Paraffin embedding was performed with controlled orientation to preserve tissue architecture. FFPE blocks were stored at room temperature until sectioning.

### Tissue sectioning and mounting

Polyethylene naphthalate (PEN) membrane slides (2 µm membrane thickness, MicroDissect GmbH) were pre-treated by UV exposure at 254 nm for 60 minutes to improve tissue adherence. Following UV treatment, sequential washing steps were performed: first in 350 mL acetone, then in a solution of 7 mL VECTABOND reagent (Vector Laboratories; SP-1800-7) diluted in 350 mL acetone, followed by a 30-second rinse in ddH2O. FFPE tissue blocks were sectioned at 3 µm thickness using a rotary microtome (Leica RM2245). Sections were mounted onto the prepared PEN membrane slides and dried at 50–56 °C prior to staining.

### Histology staining and assessment

Routine histological evaluation was performed using hematoxylin and eosin (H&E) staining. Sections were deparaffinized, rehydrated through graded ethanol, stained with hematoxylin, counterstained with eosin, dehydrated and cleared.

H&E-stained sections were assessed to confirm tissue integrity, morphology, and preservation quality. Representative sections were selected for downstream analyses based on staining quality and absence of processing artifacts.

### Immunofluorescence staining and imaging

Following tissue sectioning and mounting, slides were baked at 55°C for 20 minutes in a small oven. Tissue sections were deparaffinized and rehydrated through a graded series of xylene and ethanol washes: 2×3 minutes in xylene, 2×1 minute in 100% ethanol, 2×1 minute in 95% ethanol, 2×1 minute in 75% ethanol, 2×1 minute in 30% ethanol, and 2×1 minute in distilled water. For antigen retrieval, slides were incubated in a glycerol-supplemented Antigen retrieval buffer (1x DAKO pH9 S2367 + 10% Glycerol; water bath) at 88°C for 30 minutes, followed by a 10-minute cooldown period at room temperature. Slides were washed 2×5 minutes in distilled water and blocked with 5% BSA/ PBS for 30 min at room temperature. Primary antibodies were diluted in Renoir Red Diluent (901-PD904-052623) and incubated for 90 minutes at room temperature. Following 2×5 minutes washes in PBS, slides were incubated with secondary antibodies for 30 minutes at room temperature in 1% BSA/ PBS. All primary and secondary antibodies and respective dilutions used in this study were reported in **Suppl. Table S4**. After additional 2×5 minutes PBS washes, slides were briefly dipped in distilled water prior to nuclear counterstaining with SYTOX Green Nucleic Acid Stain (Invitrogen 57020; 1:800 dilution in distilled water) for 10 minutes. Slides were washed 2×5 minutes in PBS and mounted with Slowfade Diamond Antifade Mountant (Invitrogen, S36963). Immunofluorescence-labeled tissue samples were imaged using widefield microscopy (Zeiss Axioscan 7) equipped with a 20× dry objective. Whole tissue sections were captured with 10% tile overlap to ensure complete coverage. Fluorescent channels at 488 nm, 555 nm, 647 nm, and 750 nm were used depending on the specific antibody panel for each tissue (**Suppl. Table S4**). Illumination intensity for each channel was optimized individually. Acquired tiles were stitched together into single whole-slide images using ZEN Desk software with default settings.

### Cell type identification and segmentation

The resulting CZI files were imported individually into BIAS (Biological Image Analysis Software, version WIP 1.5.0) using the integrated import tool. The images were retiled with 1024 × 1024 pixel tiles at 5% overlap and tiles covering the respective regions of interest were selected. Cell segmentation was performed using primarily two approaches depending on cell type morphology and marker expression patterns. For well-defined cellular structures, custom-trained Cellpose models were employed for automated segmentation (Cellpose v2.0) (*58*). For cells with irregular morphologies or requiring specific marker-based identification, intensity thresholding was applied in BIAS. In rare cases where automated approaches were insufficient, manual segmentation was performed. Following segmentation, many cell types required additional classification steps including intensity-based clustering, machine learning-based classification or basic filtering based on additional marker channels. Those additional customized steps are listed in (**Suppl. Table S5**). For fallopian tube and skin cell types, CZI files were converted to SpatialData (*59*) objects using dvp-io (*60*) (version 0.1.0). Cell segmentation was performed with custom-trained Cellpose v3 (*61*) models, using Harpy (*62*) (v0.2.0) for tile-wise whole-slide processing and feature extraction. Segmentation artifacts were removed by applying manually defined area filters (26.8 μm²/900 px²) and channel-wise quantile filters (0.01–0.99th quantile for nucleus and cell membrane channels; 0–0.99th quantile for the marker channel). For skin samples, cells located outside the epithelial layer were excluded. Marker-positive cells were identified by fitting a two-component Gaussian mixture model to log-transformed marker channel intensities; only cells with a posterior probability greater than a confidence cutoff (fallopian tube: 0.95, skin: 0.90) were retained. Identified cell shapes were dilated by 1-3 μm to ensure complete cell capture, duplicates were removed, and cell area features were extracted. Three reference points were defined based on distinctive morphological characteristics on each slide to ensure precise alignment during laser microdissection and final contours were exported as .xml files. The contours were subsequently simplified by removing at least 90% of the data points defining the contour’s shape.

### Laser microdissection

Following image alignment using three manually selected tissue reference points, cell contours were imported and isolated using a Leica LMD7 system and LMD v8.5 software. The system was operated in a semi-automated manner and parameters were optimized for each tissue and cell type. The general settings can be summarized as: 63× objective, power 40-59, aperture 1, speed 5-40, middle pulse count 1, final pulse 0, offset 100, head current 45-58%, and pulse frequency 2,900. The system was aligned to each collection plate and calibrated to account for gravitational stage shift when collecting into low-binding 384-well plates (Eppendorf 0030129547). Individual cells were sorted into alternating wells with outer rows and columns left empty to prevent cross-contamination. A protective shield plate was positioned above the sample stage to minimize collection errors. For each cell type sample, a total area of 100,000 um^2^ of individually isolated cells was pooled. For each cell type, five replicas were collected, when possible, from either the same or consecutive tissue slides. Collection plates were sealed, centrifuged at 2,000g for 2 minutes, and stored at −20°C until sample processing.

### Sample preparation

1. *DVP atlas samples*

Sample processing was conducted using an Agilent Bravo automated liquid handling platform in semi-automated mode to minimize sample loss. Collection plates were removed from −20°C storage and immediately centrifuged at 2,000g for 2 minutes. Wells were washed with 28 μL of 100% acetonitrile and vacuum-dried at 45°C for 20 minutes to ensure isolated cells remained at the well bottom. Lysis buffer (6 μL of 0.013% DDM in 60 mM TEAB buffer, pH 8.5) was dispensed into each well. Plates were sealed with PCR sealing foil and heated at 95°C for 60 minutes in a PCR cycler (lid temperature 110°C). Following addition of 1 μL of 80% acetonitrile, samples were incubated at 75°C for an additional 60 minutes. After cooling, 1 μL of digestion mixture containing 4 ng/μL LysC and 6 ng/μL trypsin in 60 mM TEAB buffer was added and digested overnight (approximately 18 hours) at 37°C. The reaction was quenched with 1 μl of 10% trifluoroacetic acid. Samples were then frozen at −20°C until peptide cleanup.

ii) *Post-mortem interval tissue bulk samples*

To assess post-mortem tissue stability, an independent cohort was analyzed comprising heart and liver specimens collected across six defined post-mortem interval (PMI) time points, with ten donors per time point. Tissue collection, fixation, and processing followed the same standardized procedures as described above. To evaluate macroscopic tissue integrity and preservation over time the samples were analyzed in bulk. For bulk tissue analysis, entire tissue sections were processed from individual slides. Tissue was scraped from the slide using a scalpel, transferred to a TwinTec plate aided by 50 μL of 100% acetonitrile and vacuum-dried at 45°C until complete dryness. Samples were lysed in 42 μL of lysis buffer (70 mM TEAB, 0.013% DDM), sealed with aluminum lids, vortexed and centrifuged at 1,000g for 2 minutes. Plates were heated at 90°C for 60 minutes in a PCR cycler and allowed to cool to room temperature. Following centrifugation at 1,000g for 2 minutes, samples were sonicated using a Covaris instrument (per column: duration 300 seconds, peak power 450, duty factor 50%, cycles per burst 200, average power 225) and centrifuged at 500g for 1 minute. After addition of 7 μL of 80% acetonitrile, plates were heated at 75°C for 60 minutes and cooled to room temperature. Enzymatic digestion was performed by adding 2.5 μL each of LysC and trypsin (each 0.5 μg/μl), followed by overnight incubation at 37°C with shaking at 1,200 rpm. The reaction was quenched with 6 μL of 10% trifluoroacetic acid. Peptide cleanup was performed using SDB-RPS stage tips prepared in 200 μL pipette tips. Digested samples were centrifuged at 3,000g for 10 minutes and 20 μg peptide was loaded onto each stage tip by centrifugation at 800g for 10–15 minutes. Stage tips were washed twice with 200 μL of wash buffer 1 (isopropanol, 1% trifluoroacetic acid) and once with 200 μL of wash buffer 2 (0.2% trifluoroacetic acid) at 1,000g. Peptides were eluted with 100 μL of elution buffer (80% acetonitrile, 1.25% ammonium hydroxide) by centrifugation at 300g for 2 minutes. Eluates were vacuum-dried at 45°C for approximately 60 minutes and resuspended in 21 μL of buffer A (0.1% formic acid) with shaking at 2,000 rpm for 10 minutes at room temperature. Peptide concentration was determined by Nanodrop measurement and concentrations were adjusted to 500 ng prior to LC-MS analysis.

iii) *Tissue bulk by SPEC protocol*

For SPEC-based (*63*) bulk tissue analysis, FFPE tissue sections were scraped from slides using a scalpel and transferred to 0.5 mL tubes. Tissue was deparaffinized by three sequential incubations in 100 μL of n-heptane at 30°C for 1 minute with shaking at 800 rpm, discarding the solution after each step. Samples were then washed once with 100 μL of methanol under identical conditions. Deparaffinized tissue was lysed in 100 μL of lysis buffer (50 mM TEAB, 40 mM CAA, 10 mM TCEP, 2% SDC) at 98°C for 10 minutes. Following cooling on ice, samples were sonicated using a Branson sonicator (setting 2, duty cycle 20%) for 3 minutes or until homogeneous and clear. Homogenized lysates were heated again at 98°C for 10 minutes, cooled on ice and protein concentration was determined by tryptophan assay. Samples were adjusted to 1 μg/μL concentration. For on-tip digestion, strong anion exchange (SAX) stage tips (double layer) were primed with 10 μL of DMSO and centrifuged at 700g for 1.5 minutes, followed by equilibration with 20 μL of equilibration buffer (20 mM CAPS pH 10.5, 0.01% DDM) at 700g for 2 minutes. 200 ng Protein lysate was diluted 1:10 in loading buffer (20 mM CAPS, 50% methanol, 0.01% DDM) and loaded onto the tip by centrifugation at 400g for 15 minutes. Tips were washed with 20 μL of washing buffer (50 mM TEAB, 0.01% DDM) at 700g for 1.5 minutes. Digestion buffer (5 μL: 50 mM TEAB, 10 mM TCEP, 0.01% DDM, 0.05 μg/μL LysC and trypsin) was added, tips were pulse-spun at 100g for 20 seconds and samples were incubated at 37°C for 1 hour.

### Peptide loading

Peptide purification was performed using C18 Evotips (Evotip Pure, EvoSep). Sample plates were thawed, centrifuged at 2,000g for 2 minutes, and maintained on ice. Evotips were activated in 1-propanol for 3 minutes, washed twice with 50 μL of buffer B (0.1% formic acid, 99.9% acetonitrile), and centrifuged at 700g for 1 minute between washes. Following re-activation in 1-propanol for 3 minutes, tips were equilibrated with two 50 μL of buffer A (0.1% formic acid) washes. Sample loading was performed as follows:

1. For DVP samples was performed by adding 70 μL of buffer A per tip followed by sample transfer. Each well was rinsed with 10 ìL of buffer A and combined with the corresponding tip. Peptide binding was achieved by centrifugation at 700g for 2 minutes.
2. For post-mortem interval tissue bulk samples, 500 ng in 50 μL of buffer A was added per tip and centrifuged at 700g for 2 minutes.
3. For SPEC-processed bulk samples, peptides were eluted directly from SAX tips into prepared Evotips using 20 μL of elution buffer (1% formic acid) by centrifugation at 300g for 5 minutes.

Tips were washed with 50 μL of buffer A, overlaid with 150 μL of buffer A and centrifuged at 700g for 15 seconds. Loaded tips were stored in Evotip boxes with fresh buffer A for a maximum of 3 days prior to LC-MS analysis.

### LC-MS/MS analysis

1. *DVP atlas samples & Tissue bulk by SPEC protocol*

The samples were analyzed using an Evosep One liquid chromatography system coupled to an Orbitrap Astral Zoom mass spectrometer (Thermo Fisher Scientific) via an EASY-Spray source operating at 1,900 V. Peptides were separated on an Aurora Rapid C18 column (5 cm × 75 μm ID, 1.7 μm particle size, IonOpticks) maintained at 60°C using a Whisper Zoom 80 SPD gradient. Data-independent acquisition was performed with Orbitrap MS1 scans recorded at 240,000 resolution over a scan range of 380–980 m/z using 500% normalized AGC target and 3 ms maximum injection time.

For bulk SPEC samples, a total of 200 equidistant isolation windows with 3 Th width were defined covering the precursor selection range of 380–980 m/z (**Suppl. Table S6**). Fragment ion spectra were acquired with 6 ms maximum injection time, 500% normalized AGC target and 25% HCD collision energy.

For DVP samples, the Orbitrap Astral was equipped with a FAIMS Pro interface (Thermo Fisher Scientific) operated at −40 V compensation voltage with 3.5 L/min carrier gas. Isolation window width was optimized according to precursor density, with 100 variable-width windows covering the precursor selection range of 380–980 m/z based on the precursor distribution of a reference sample using pyDIAid (*64*) (**Suppl. Table S7**). Fragment ion spectra were acquired with 10 ms maximum injection time, 500% normalized AGC target and 25% HCD collision energy.

1. ii) *Post-mortem interval tissue bulk samples*

Samples were analyzed using an Evosep One liquid chromatography system coupled to a timsTOF HT mass spectrometer (Bruker Daltonik) via a nanoelectrospray ion source. Peptides were separated on a 15 cm × 75 μm column packed with 1.9 μm C18 beads (Evosep) using a 44-minute gradient (30 SPD) at a flow rate of 250 nL/min. Mobile phases consisted of 0.1% formic acid in water (A) and 0.1% formic acid in acetonitrile (B). A 10 μm ID zero dead volume electrospray emitter (Bruker Daltonik) was used for ionization. Data-independent acquisition was performed with 20 isolation windows spanning an m/z range of 350–1,200 and an ion mobility range of 0.7–1.3 Vs·cm⁻² at a cycle time of 2.2 seconds. Collision energy was ramped linearly as a function of ion mobility from 20 eV at 1/K0 = 0.6 Vs·cm⁻² to 59 eV at 1/K0 = 1.6 Vs·cm⁻².

### Raw data processing

1. *DVP atlas samples & Tissue bulk by SPEC protocol*

DVP and bulk tissue samples were treated as individual entities. All raw files were analyzed using DIA-NN (version 1.9.2) in library-free mode. A spectral library was predicted by deep learning from a human reference proteome FASTA file from UniProt (UP000005640, taxon ID 9606, reviewed entries only, downloaded March 22, 2024). Search parameters included: trypsin/P digestion with maximum one missed cleavage, N-term M excision enabled, peptide length range of 7-30 amino acids, precursor charge range of 1-4, precursor m/z range of 300-1,800, and fragment m/z range of 200-1,800. Mass accuracy, MS1 accuracy and scan window were set to 0.0 (automatic). Heuristic protein inference was enabled with protein inference from genes. The neural network classifier was set to single-pass mode, quantification strategy to ’Quant UMS (high precision)’, cross-run normalization to ’RT-dependent’, and library generation to ’IDs, RT & IM profiling’. The generated spectral library was used for all cell type analyses. Individual replicates from all cell types/ tissues were searched together in one search with MBR and peptidoforms enabled. The resulting 27 report files were combined and protein intensities were normalized using directLFQ (version 0.2.19) with standard settings. No missing value imputation was applied.

ii) *Post-mortem interval tissue bulk samples*

Bulk post-mortem samples were processed using DIA-NN (version 1.8.1) with identical search parameters except for fixed mass accuracy settings of 9 ppm (MS2) and 16 ppm (MS1) and a scan window radius of 7. Replicates were normalized using directLFQ (version 0.2.19) with standard settings.

### Data analysis

Data analysis and visualization were performed in Python (version 3.12.11) using NumPy (v2.3.3), pandas (v2.3.3), seaborn (v0.13.2), matplotlib (v3.10.6) and scikit-learn (v1.7.2). Overrepresentation and gene set enrichment analyses were performed using gseapy (v1.1.10).

1. *Data filtering and preparation*

Protein intensities from the directLFQ output file ‘report.tsv.protein_intensities.tsv’ were used for all subsequent analyses. To evaluate the effect of normalization, log₁₀ median protein intensity distributions were compared across cell types before and after directLFQ normalization. For each cell type, five biological replicates were collected where possible. When reporting cell type-level measurements or identification, median protein intensities were calculated across replicates, with values set to missing if more than half of the replicates lacked a measurement for a given protein. For MS-based analysis, protein detection was defined as a non-missing intensity value.

ii) *Statistical analysis and data visualization*

To assess proteome stability across post-mortem intervals, the number of identified protein groups was determined per sample across six timepoints (12, 18, 24, 36, 48 and 72 hours). For the ranked protein intensity distribution, the maximum intensity across all cell types was calculated for each protein, log₁₀-transformed, and proteins were ranked in descending order. Protein class detection rates were assessed using protein class annotations from the Human Protein Atlas. For each class, the proportion of annotated proteins detected in the dataset was calculated. To identify proteins lacking prior experimental evidence at the protein level, reviewed human protein-coding sequences were obtained from UniProt, stratified by protein existence level (PE1–PE5). Each list was matched to the dataset by UniProt accession. Proteins with PE2 (transcript evidence), PE3 (homology inference), PE4 (predicted) or PE5 (uncertain existence) classifications that were detected in at least one cell type were reported. For each protein, (proteotypic-) peptides were manually inspected using Skyline-daily version 26.0.9.032 (*65*, *66*). Cell type specific spectral libraries were constructed from DIA-NN search results (details under *Raw data processing*) and corresponding raw files were loaded to visually assess individual extracted ion chromatograms. Selected peptides were visualized using alphadia-validate (*67*). To characterize proteins absent from the dataset, identified proteins were compared to the complete human reference proteome (UniProt UP000005640). Tissue enrichment analysis of absent proteins was performed using the GTEx database (GTEx_Tissues_V8_2023; terms differing only by age or sex were consolidated; adjusted p-value < 0.5). To assess functional annotation coverage, multiple Enrichr libraries were screened and the three with highest coverage were selected: InterPro_Domains_2019, GO_Molecular_Function_2023 and GO_Biological_Process_2023 (adjusted p-value < 0.05). The proportion of absent proteins explained by tissue-specific expression, functional annotation, both or neither category was summarized. The number of unique protein identifications was determined per cell type across individual replicates. Coefficient of variation was calculated across replicates for each protein per cell type to assess reproducibility. Per-cell-type intensity profiles were generated by ranking proteins by median intensity within each cell type. For marker protein visualization, relative expression was calculated as the proportion of total expression across all cell types and absolute expression was represented by min-max normalized intensity values. Within-tissue marker validation was performed using established cell type markers, most of which were used for immunofluorescence-guided cell identification in the DVP pipeline, comparing raw protein intensity across cell types isolated from the same tissue. To assess overlap between testis-specific proteins and the oocyte proteome, a list of proteins annotated as detected exclusively in testis was obtained from the Human Protein Atlas and compared to proteins identified in oocytes by UniProt accession. The core proteome comprised proteins detected across all 27 cell types; cell type-specific proteins were those detected in a single cell type. Overrepresentation analysis of core proteome and cell type-specific proteins was performed against the KEGG_2021_Human database (adjusted p-value < 0.05). Principal component analysis was performed on the core proteome using scikit-learn. Log₂-transformed protein intensities were standardized using StandardScaler prior to PCA. Analysis was performed on individual replicates. Gene set enrichment analysis was performed for each cell type. Proteins were ranked by log₂ fold change relative to background, calculated as the median intensity across all cell types. GSEA was run against KEGG_2021_Human with minimum gene set size of 5 and maximum of 500 (adjusted p-value < 0.05). Disease-related pathways were excluded by keyword filtering. The 20 pathways with greatest variance in normalized enrichment score across cell types were selected for visualization. Pathway proportion was calculated as the sum of intensities of pathway member proteins divided by total protein intensity per cell type. Intersection analysis of shared proteins across cell types was performed based on binary protein detection per cell type, showing the top 20 intersections ranked by size.

iii) *RNA – protein comparison*

For data integration and to enable direct comparison between protein-centric and transcript-centric datasets, UniProt identifiers were mapped to gene-level identifiers from the Human Protein Atlas (HPA), based on Ensembl v109 annotations and Uniprot release 2022_05. This mapping resulted in the exclusion of a subset of proteins due to mismatched or missing identifier overlap. To further ensure robust comparisons and minimize false-positive assignment, protein groups (i.e. proteins that could not be uniquely mapped to a single gene) were classified as multi-mappable and excluded from downstream analysis (195 protein groups). Following filtering, the final dataset comprises 19297 genes (Ensembl v 109). The single cell type RNA expression profiles presented in the HPA Single Cell resource are based on publicly available scRNAseq and snRNAseq datasets, representing tissues of the human body, mapped and annotated by the HPA. Original references and more details for each RNAseq dataset are listed in **Suppl. Table S8**. Details of the data processing can be found in the HPA method details (*68*). To enable comparison with proteomics data, relevant single cell RNAseq clusters were selected for each tissue based on the best possible overlap to the DVP sampling. Cluster selection was performed independently for each tissue, and the full list of included clusters is provided in **Suppl. Table S2**, along with the number of cells, cell type annotations, and information on whether the data is based on single-cell or single-nuclei RNAseq. For most cell types (e.g. hepatocytes, oocytes, granulosa, and immune cell populations), cluster selection was straightforward. However, for anatomically complex structures represented in DVP samples, multiple RNAseq clusters were required. For example, renal glomeruli include both podocytes and vascular endothelial cells, therefore, cluster c22 (podocytes) and c23 (glomerular capillary endothelial cells) were jointly used to represent this structure. When multiple clusters were selected, either within the same cell type or mix of different cell types, a weighted mean expression value was calculated to generate a pseudo-bulk profile. To validate cluster selection, gene expression profiles across all clusters available for respective tissues, were correlated with protein abundance profiles from corresponding DVP samples within the same tissue type. Examples of this correlation analysis are shown in **Fig. S13** for liver, ovary and lymph node, remaining tissues are added to the method background page. This resulting dataframe contained two modalities (DVP and RNAseq), 27 cell types and a listing of all identified transcripts and proteins. RNA detection was defined as expression ≥ 1 nCPM whereas MS-based protein detection was defined as non-missing intensity value for all following analysis. Spearman rank correlations were computed pairwise between all RNA and MS cell types using log2-transformed expression values matched by

Ensembl gene identifier. Results were visualized as a heatmap of all pairwise Spearman *r* values. To facilitate identification of relatively higher-correlating MS cell types for each RNA cell type, a row-normalized version was computed by z-scoring each row across MS cell types. To compare gene signatures between proteomes and transcriptomes of the different cell types, we applied the R Package Ucell (*30*) (version 2.2.0). Ucell scores a given signature in a sample based on its genes ranks using Mann-Whitney U statistic. HALLMARK pathways (*69*) were used as the list of given signatures. For each cell type and each HALLMARK pathway, a Ucell score was calculated for the protein and RNA pooled dataset independently. Prior to classification and RNA comparison, six DVP cell types were consolidated (by using Max-value representation) into three broader cell type groups. While values for the individual cell types are displayed separately in the resource to ensure transparency, alveolar cell 1 and 2 were grouped into alveolar cells, ciliated and secretory cells of the fallopian were grouped into fallopian epithelial cells, and finally, CD4 and CD8 T-cells were grouped into T-cells. These groupings were applied due to a high degree of protein overlap between the respective cell types, and they facilitate a clearer overview of the data by highlighting proteins with biologically relevant cell type-enriched expression. Per-gene detection agreement scores were computed across all cell types for genes present in both the RNA and MS datasets. Detection was binarized per cell type as detected or not detected, independently for RNA and MS. The two binary vectors were combined into a four-category encoding per cell type: both detected (3), RNA only (2), MS only (1), or neither (0). The agreement score was calculated as the proportion of cell types in concordant states (both detected or both absent). An analogous expression-level agreement score was computed using the same formula, replacing the detection binary with an above/below per-gene median binarization, calculated independently for RNA and MS across all cell types. Gene expression profiles were categorized along two independent dimensions: expression distribution and expression specificity (**Suppl. Table S3**). This dual framework allowed us to capture both the distribution of expression across cell types and the relative enrichment of genes in specific cellular contexts. These criteria were applied to the RNA dataset selected for the comparison with the DVP data. As a result, gene classification may differ from those presented online for the same genes, which is based on the full dataset. The same distribution and specificity criteria were also applied to the DVP data, enabling a direct comparison at the category level between RNA expression and protein detection. Next, we compared the overlap and matched elevated cell types between RNA- and protein-based specificity categories. To simplify the comparison and improve readability, we treated all elevated categories as a single group, without distinguishing between enriched, group enriched and enhanced expression. Specificity overlap was defined as full overlap (exact match) or partial overlap (at least one overlapping cell type) combined. In addition, a small number of cell types were treated as exceptions and classified as overlaps even when the elevated protein signal was detected in a different cell type than corresponding RNA. These exceptions included: RNA detected in neurons with protein in astrocytes or microglia, RNA detected in pancreatic epithelial cells with protein detected in pancreatic islet cells, RNA detected in granulosa cells with protein detected in oocytes. These cases showed substantial overlap at protein level, and with biological and sampling reasons behind the overlap, and rather than grouping them together (such as alveolar cells and T-cells described above), we defined them as exceptions to the overlap rules. Pathway enrichment analyses were performed using Enrichr via the GSEApy interface, testing gene lists derived from each quadrant of the overlap × expression agreement score matrix against the GO Biological Process 2023 gene set library. Results were filtered to adjusted p-value < 0.05 and the top four terms per gene list were visualized as normalized bar plots, with bar length scaled relative to the most significant term within each panel.

iv) *Bulk-DVP comparison*

The IHC-based protein profiles were downloaded from HPA (normal_ihc_data), the list of 76 cell types and 45 normal tissues, covering 15312 protein-coding genes, was filtered by overlapping tissue types resulting in 18 different tissues to compare. Blood vessel tissue is missing IHC annotation, and IHC annotated small intestine includes both ileum and jejunum. The comparison was based on detection, meaning all levels of detection were viewed as positive, and IHC detection was assigned if detected in either of the cell types annotated in respective tissue types. Integration of bulk RNA sequencing data was straightforward, as both the RNA and MS datasets were generated using bulk profiling approaches. Bulk RNAseq expression profiles were downloaded from the HPA for tissue types overlapping with those represented in the MS bulk dataset. The comparison was based on the HPA consensus expression data, which integrates HPA in–house RNAseq data with expression profiles from the GTEx project (*20*). Prior to classification of expression profiles and downstream overlap analysis, tissue type grouping was applied in accordance with the HPA tissue resource. Specifically, intestinal tissues were represented by a single value, where duodenum, ileum, jejunum, and colon were grouped and assigned the maximum expression value within the tissue group. Similarly, the spleen and lymph node were grouped into lymphoid tissue. The specificity and distribution categories for the bulk RNA and MS data respectively are available in the resource for further exploration of the category overlap and agreement scores across modalities. Detection overlap across bulk MS, bulk RNAseq, DVP, and sc/snRNAseq was visualized as an UpSet plot using the upsetplot library (v.0.9.0), with genes classified as detected or not detected in each modality. Detection overlap between cell types and their respective bulk modality was visualized as Venn diagrams using the matplotlib-venn library (v.1.1.2), with genes classified as detected or not detected in each cell type. To characterize proteins uniquely detected by bulk MS, and missing in the DVP samples, the full set of snRNAseq clusters representing the kidney was utilized, including the 16 main cell types (based on 36 different sn clusters combined into weighted mean nCPM to represent each of the main cell type, more details can be found on the atlas method pages). For each matched gene to protein, the maximum nCPM across the different cell type clusters was noted and summarized based on frequency. Pathway enrichment analyses were performed using Enrichr via the GSEApy interface, testing gene lists corresponding to selected UpSet plot intersections against the KEGG 2021 Human gene set library. Disease- and infection-related terms were excluded prior to visualization. Results were filtered to adjusted p-value < 0.05 and the top four terms per gene list were visualized as normalized bar plots, with bar length scaled relative to the most significant term within each panel. Lists of known CTAs were extracted from the CTDatabase further filtered and extended by *Djureinovic et al.* (*32*) (**Suppl. Table S1 and S5).** CTA expression in oocytes was manually inspected similar to the process used for protein evidence evaluation (see above). In addition, precursor evidence in oocytes was contrasted to the respective absence in corresponding bulk ovary dataset. Both single oocytes as well as bulk ovary raw files were imported into Skyline-daily version 26.0.9.032 (*64*, *65*) with an oocyte specific spectral library with retention time filtering set to include matching scans within 2 min of the MS/MS identification. Raw times and raw intensities of individual fragment ions of oocyte specific CTAs were exported and fragment ion traces were plotted using python. A window of 2 min around each CTA identification in oocytes was inspected for the bulk ovary dataset to exclude retention time shifts as a potential explanation for their respective absence in a side by side comparison. A condensed window of ∼0.25 min was used to increase readability in respective figures.

### IHC stainings

All Immunohistochemical (IHC) images with chromogen detection (i.e. not fluorescent images used for cell type identification) are from the HPA, freely available online, and listed with gene target, antibody ID and direct link to more antibody information in **Suppl. Table S9**.

### Data integration into HPA

The proteomics data, including both DVP and bulk MS datasets, were integrated into version 25 of the HPA. The DVP data, providing cell type resolved measurements, were incorporated into the single cell resource, whereas the bulk MS data was integrated into the tissue resource. This structure enables protein detection data to be presented alongside corresponding RNAseq data at matching levels of resolution. Protein intensity values are visualized as interactive bar plots, where clicking individual bars, or corresponding cell type or tissue labels, provides access to intensity values for each replicate. HPA-defined categories, including specificity and distribution (**Suppl. Table S3**), are available as filters to facilitate user friendly exploration of the dataset. In addition, the total number of proteins within each category per cell type or tissue is displayed on the respective summary pages. In addition, comprehensive method pages are provided to describe data generation and processing. These pages overlap with the present Methods section but include expanded explanations, examples, and technical details enabled by the dedicated portal. Furthermore, each cell type and tissue are described in curated knowledge-based summary pages. Agreement scores and specificity overlaps are integrated into the resource and can be used as filters in combination with other search criteria. For DVP datasets, representative fluorescence images used for shape definition and sample collection are included. Additionally, RNA correlation heatmaps are provided for tissues relevant to DVP comparisons and are accessible via the detailed method pages.

## Data availability

Imaging data have been deposited to BioImages (*57*) with the accession number S-BIAD2869. The complete data set is integrated into the HPA Tissue and Single Cell Resource creating a freely accessible, multi-modal reference enabling community-wide exploration across MS, RNA and antibody-based modalities (v25.proteinatlas.org/humanproteome/single+cell/dvp). The mass spectrometry proteomics data, Supplementary Tables and generated code will be available upon publication.

---

**Fig. S1.**
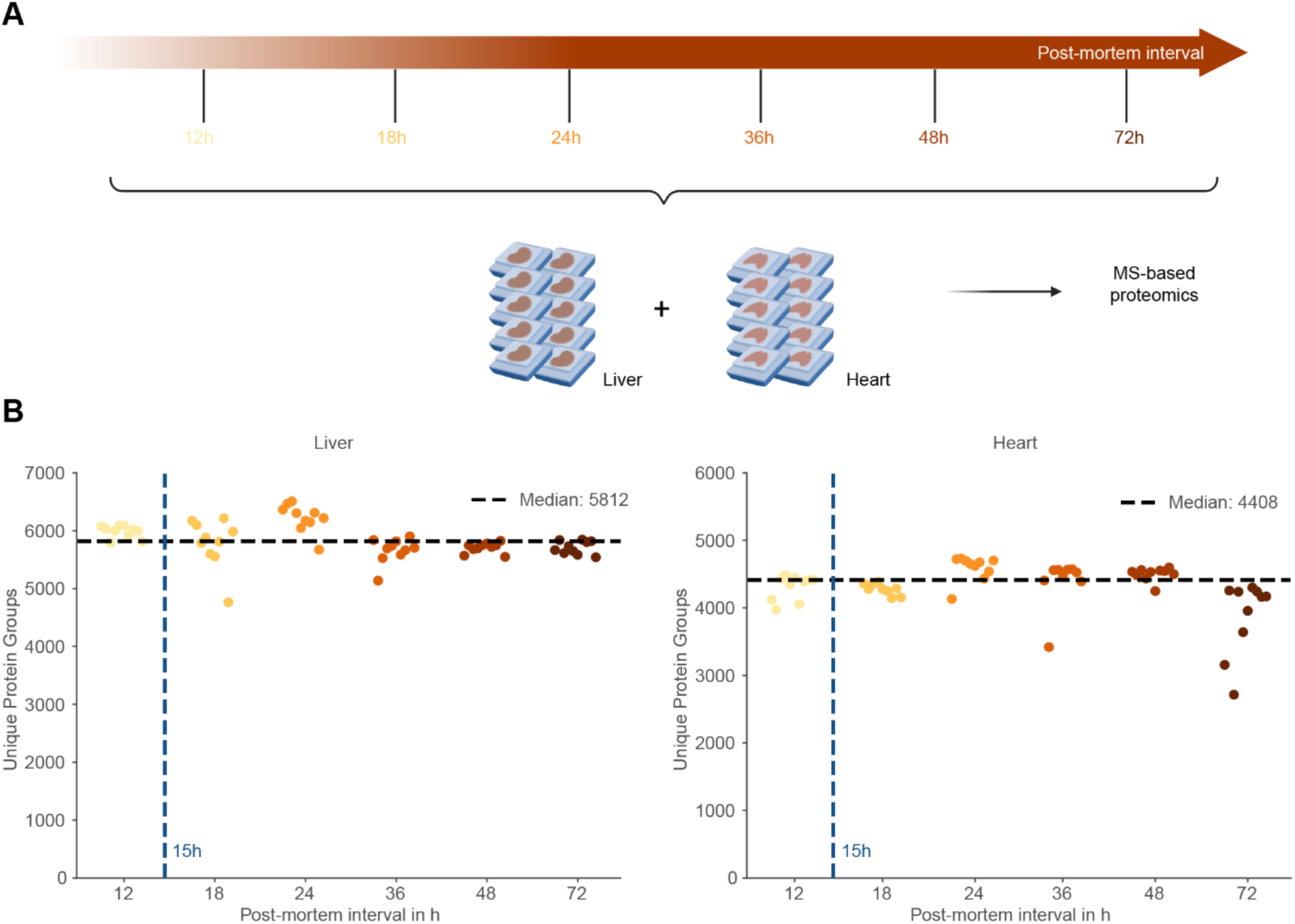
Proteome depth remains stable across extended post-mortem intervals. (**A**) Schematic of experimental design. Formalin-fixed paraffin-embedded liver and heart tissue sections with defined post-mortem intervals (12, 18, 24, 36, 48, and 72 hours) were analyzed by bulk mass spectrometry-based proteomics. (**B**) Number of unique protein groups identified per sample across post-mortem intervals in liver (left) and heart (right). N = 10 samples per time point. Overall median indicated by a black dotted line. Blue dotted line shows PMI of the samples used in the study.

**Fig. S2.**
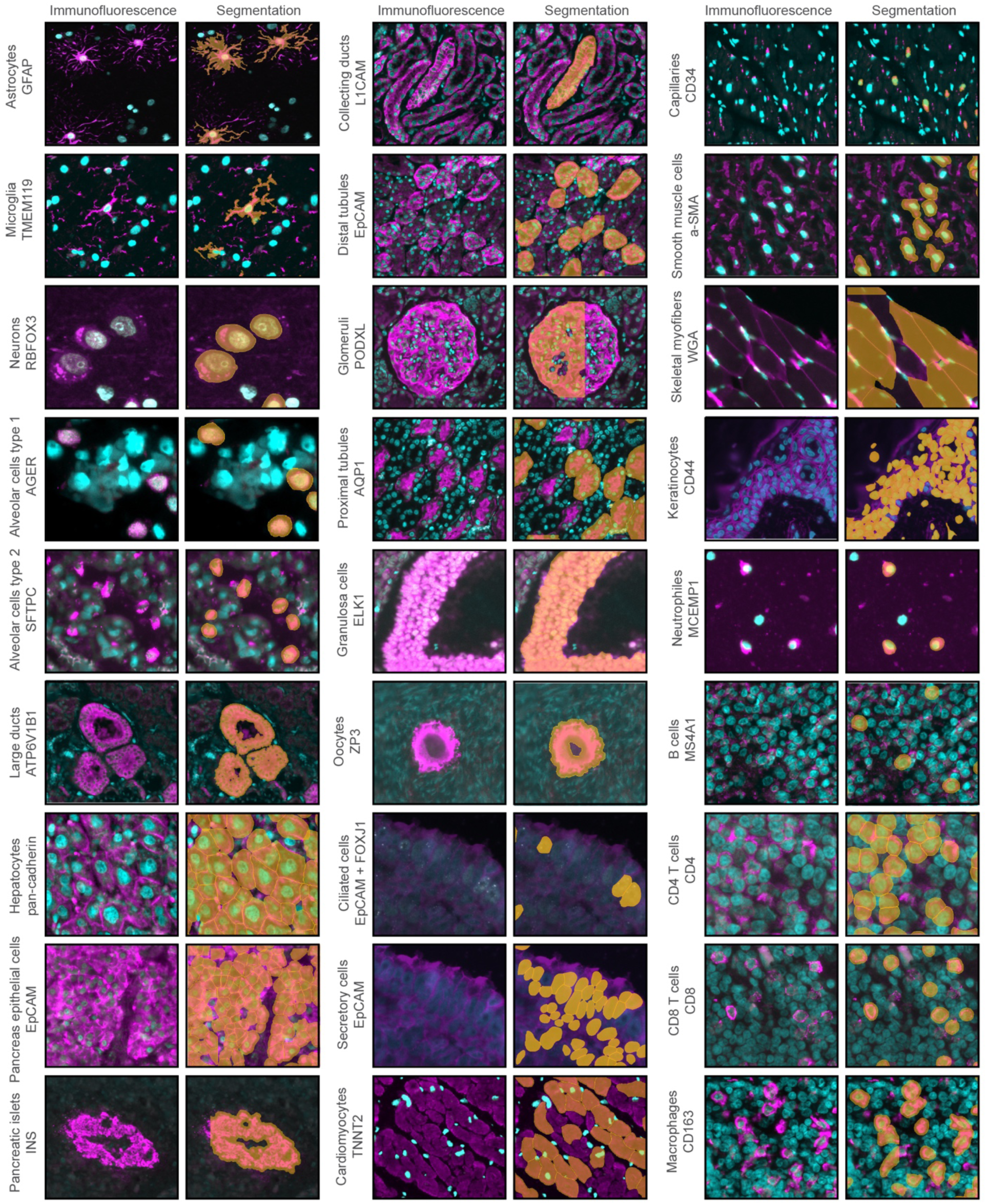
Identification and segmentation of 27 cell types. Representative immunofluorescence images (left) and corresponding segmentation masks (right) for all cell types analyzed in this study. Cell type and marker protein are indicated for each row. Nuclei are shown in cyan, cell-type-specific markers in magenta, and segmentation contours in orange.

**Fig. S3.**
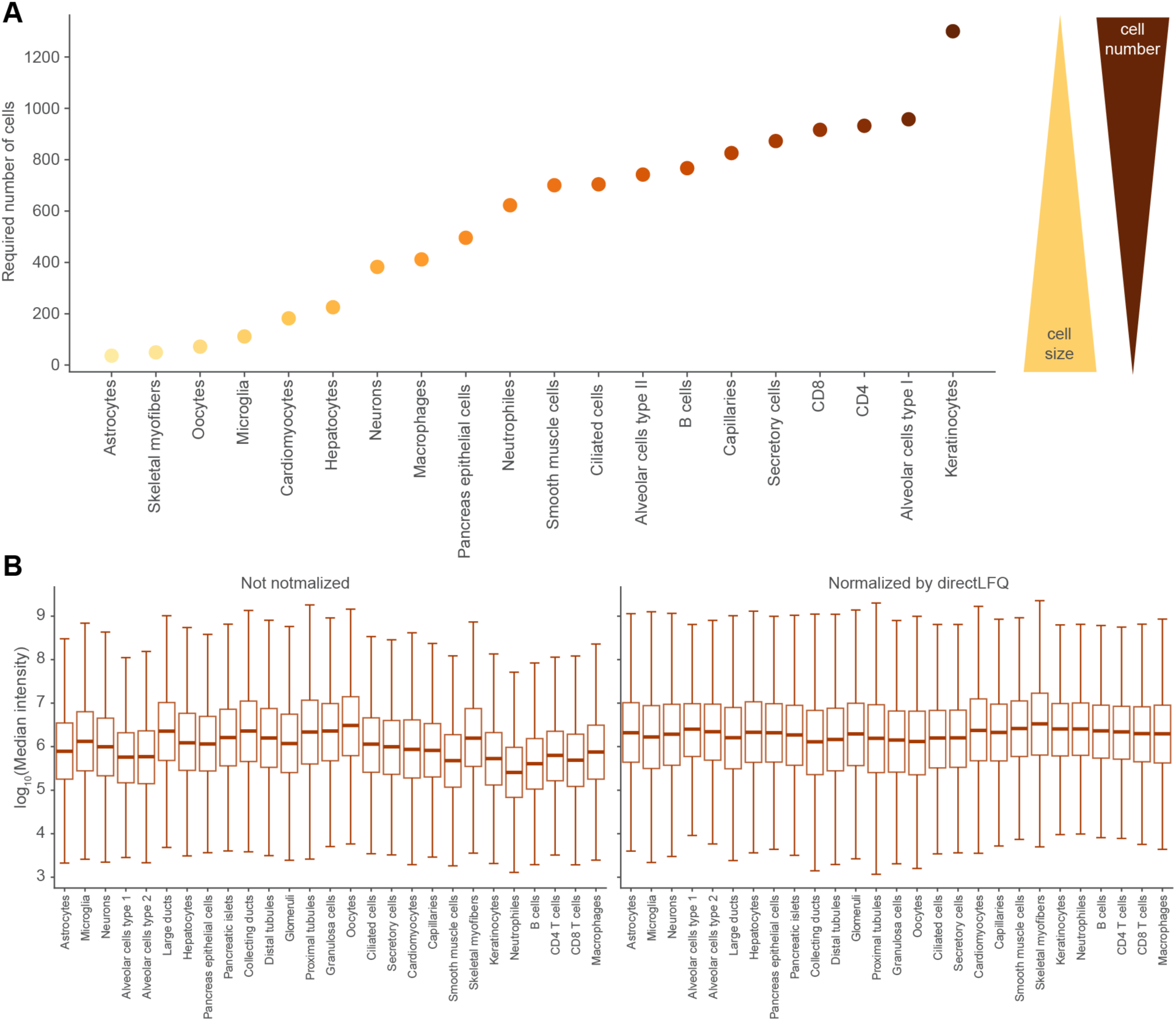
Area-based cell collection enables protein intensity normalization across cell types. **(A)** Required number of cell contours per cell type to reach a cumulative area of 100,000 μm². Cell types are ordered by increasing cell count. Inset illustrates the inverse relationship between cell size and required cell number. **(B)** Distribution of log₁₀ median protein intensity per cell type before (left) and after (right) normalization using directLFQ.

**Fig. S4.**
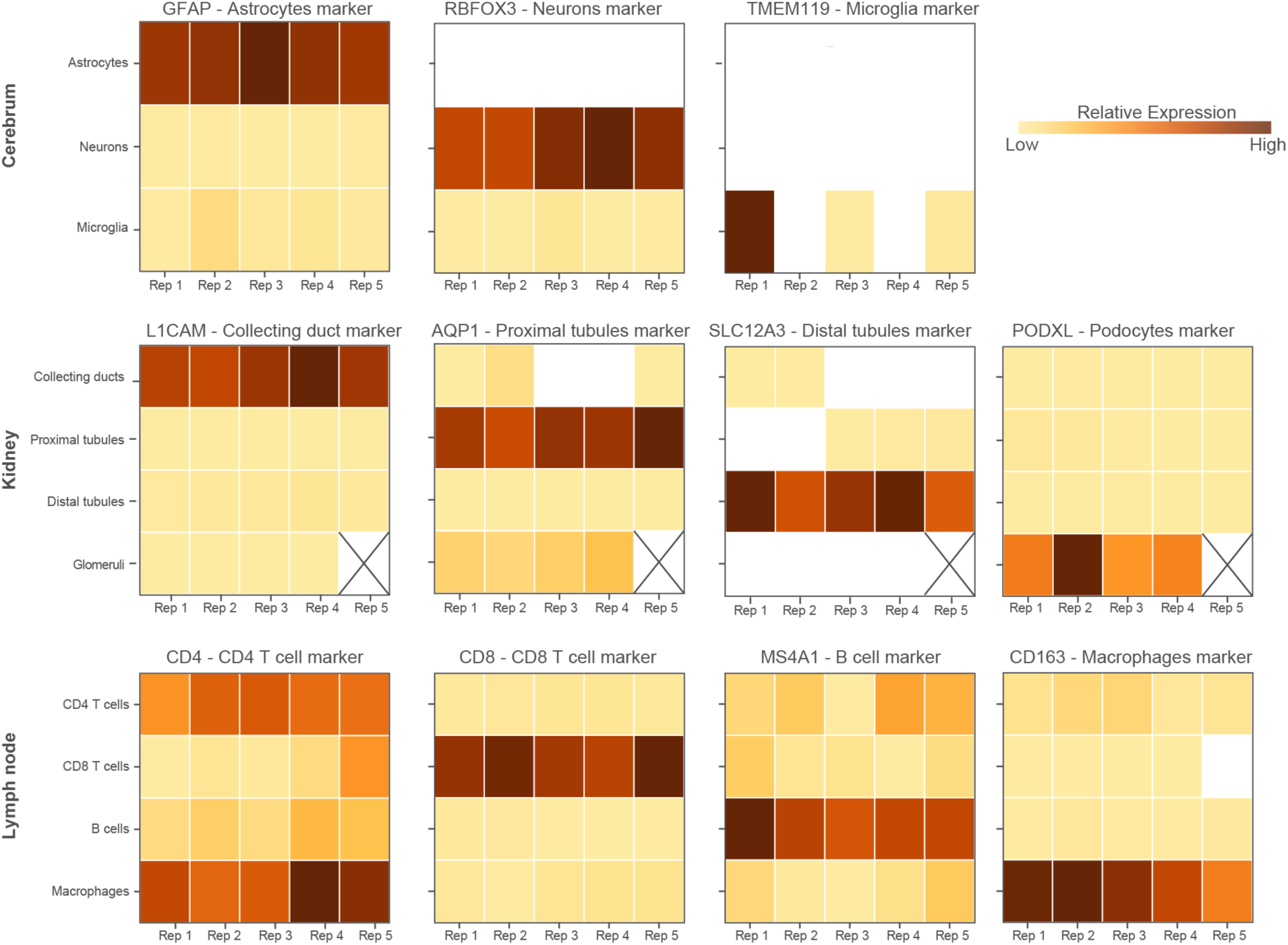
Cell-type-enriched proteomes confirmed by distinct marker protein profiles within shared tissues. Heatmaps displaying protein intensity of established cell-type markers (indicated above each heatmap) across cell types isolated from the same tissue. Cerebrum (top): astrocytes, neurons and microglia. Kidney (middle): collecting ducts, proximal tubules, distal tubules and podocytes. Lymph node (bottom): CD4 T cells, CD8 T cells, B cells and macrophages. Individual replicates are shown. White boxes indicate no detected signal and crossed boxes indicate missing replicates.

**Fig. S5.**
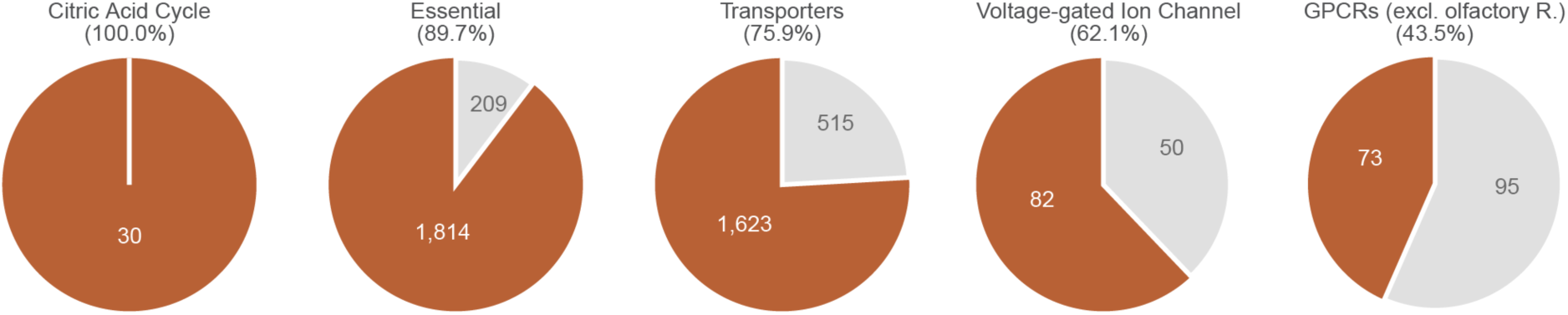
Extended protein class coverage analysis. Pie charts showing the proportion of detected (orange) versus undetected (grey) proteins for five protein classes annotated by the Human Protein Atlas (additional to Fig. 2B). Percentages indicate detection rates. Numbers show absolute protein counts.

**Fig. S6.**
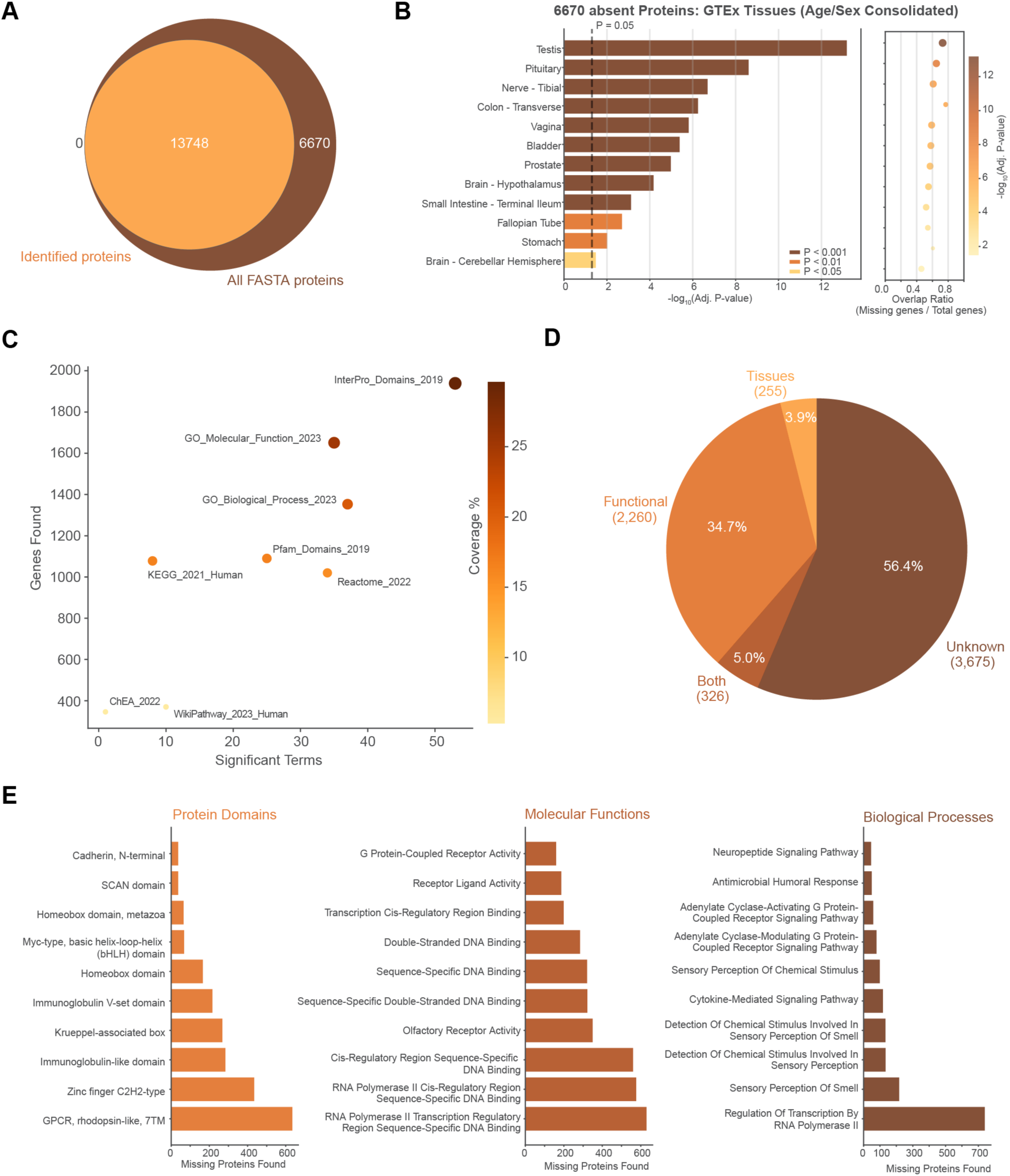
Absent proteins are explained by tissue specificity and functions beyond the analyzed cell types. (**A**) Venn diagram comparing proteins identified in this study to the complete human FASTA reference proteome (UP000005640). (**B**) Tissue enrichment analysis of absent proteins using the GTEx database (terms differing only by age or sex were consolidated). Bar length indicates -log₁₀(adjusted P-value). Dot plot (right) shows overlap ratio of missing genes to total pathway genes. (**C**) Comparison of enrichment libraries by coverage of absent proteins. Color indicates percentage of absent proteins annotated. (**D**) Pie chart summarizing the proportion of absent proteins explained by tissue-specific expression, functional annotation, both, or unknown. (**E**) Top enriched terms among absent proteins for protein domains (left), molecular functions (center), and biological processes (right). Bar length indicates the number of absent proteins per term.

**Fig. S7.**
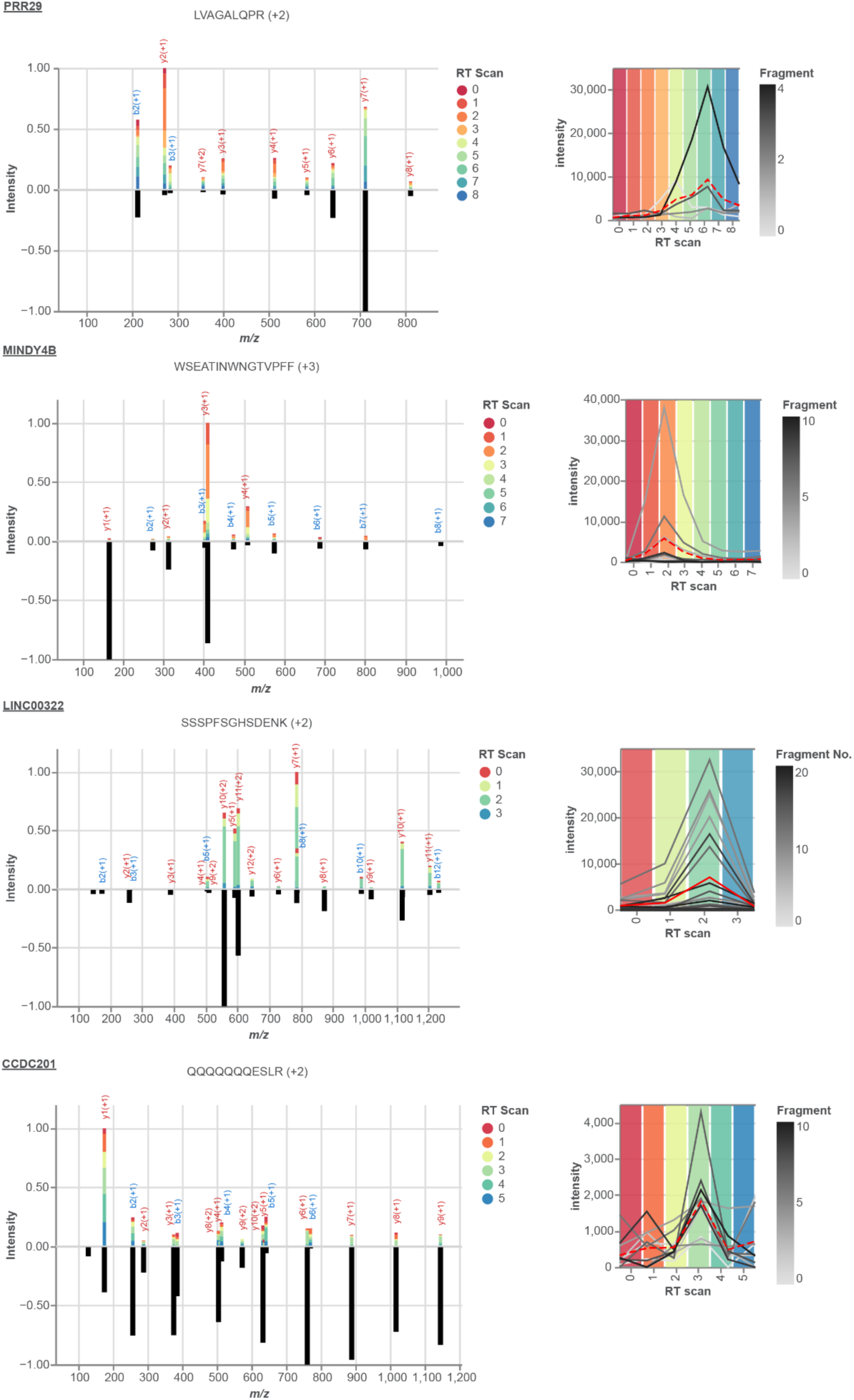
Validation of detected proteins of protein existence (PE) groups below PE1. Spectral evidence for selected proteotypic peptides of PRR29 (PE2), MINDY4B (PE3), LINC00322 (PE4) and CCDC201 (PE5). The left panel shows a mirror plot. The top is a stacked bar representation of matched fragment ions across consecutive scans (RT Scan: retention time scan). The bottom half represents a matched predicted fragment spectrum. The right panel shows an extracted ion chromatogram of matched fragments (grey) and their average peak representation (red). The color-coded RT Scans between the left and right panel of each peptide are matched.

**Fig. S8.**
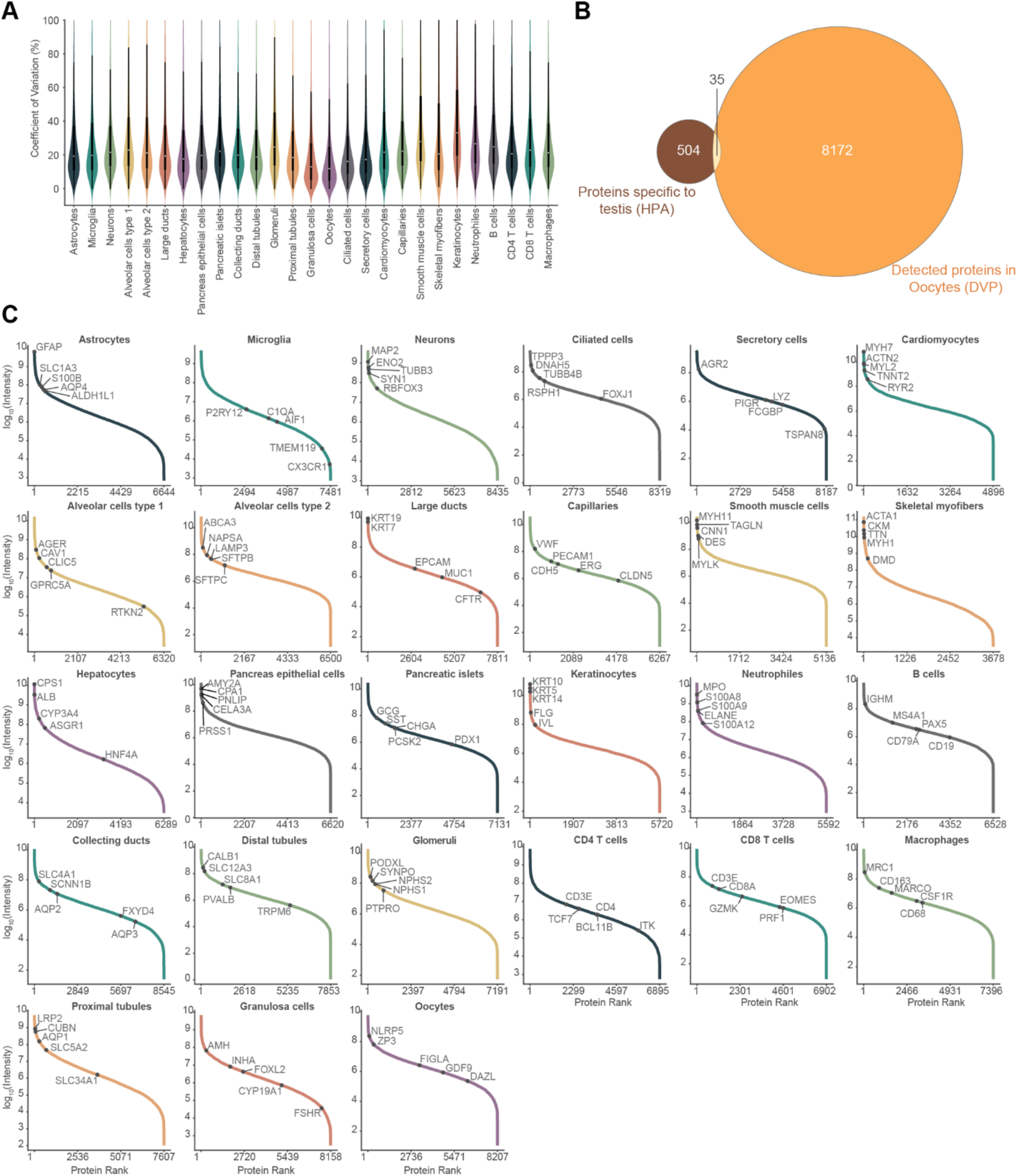
Technical reproducibility and cell type-resolved proteome characteristics. (**A**) Distribution of coefficient of variation (CV) across all identified proteins per cell type. (**B**) Venn diagram comparing proteins annotated as testis-specific by the Human Protein Atlas with proteins detected in oocytes by DVP. (**C**) Ranked protein intensity distribution per cell type. Proteins are ordered by log₁₀ median intensity. Selected cell type-characteristic markers are labeled. Protein rank range is indicated on the x-axis.

**Fig. S9.**
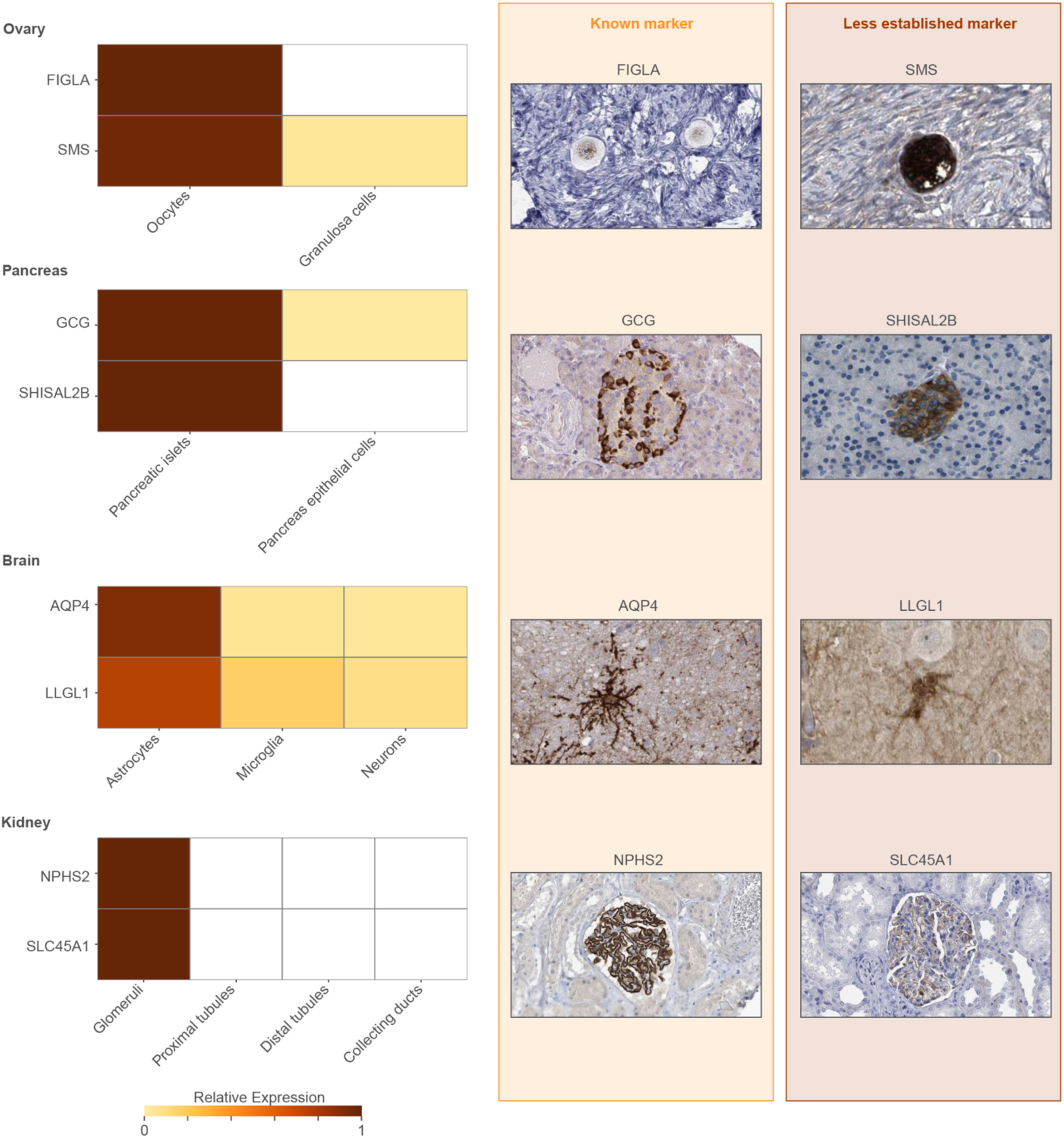
Known and less established cell type markers validated by relative expression and immunohistochemistry. Heatmaps (left) show relative protein expression of selected known and less established markers across cell types for four tissues: ovary (oocytes, granulosa cells), pancreas (pancreatic islets, pancreatic epithelial cells), brain (astrocytes, microglia, neurons), and kidney (glomeruli, proximal tubules, distal tubules, collecting ducts). Color indicates relative expression level (0–1 scale). Corresponding immunohistochemistry images from the HPA (antibody details listed in Suppl. Table S9) are shown for known markers (center, orange): FIGLA (oocytes), GCG (pancreatic islets), AQP4 (astrocytes), and NPHS2 (glomeruli), and for less established marker candidates (right, brown): SMS (oocytes), SHISAL2B (pancreatic islets), LLGL1 (astrocytes), and SLC45A1 (glomeruli).

**Fig. S10.**
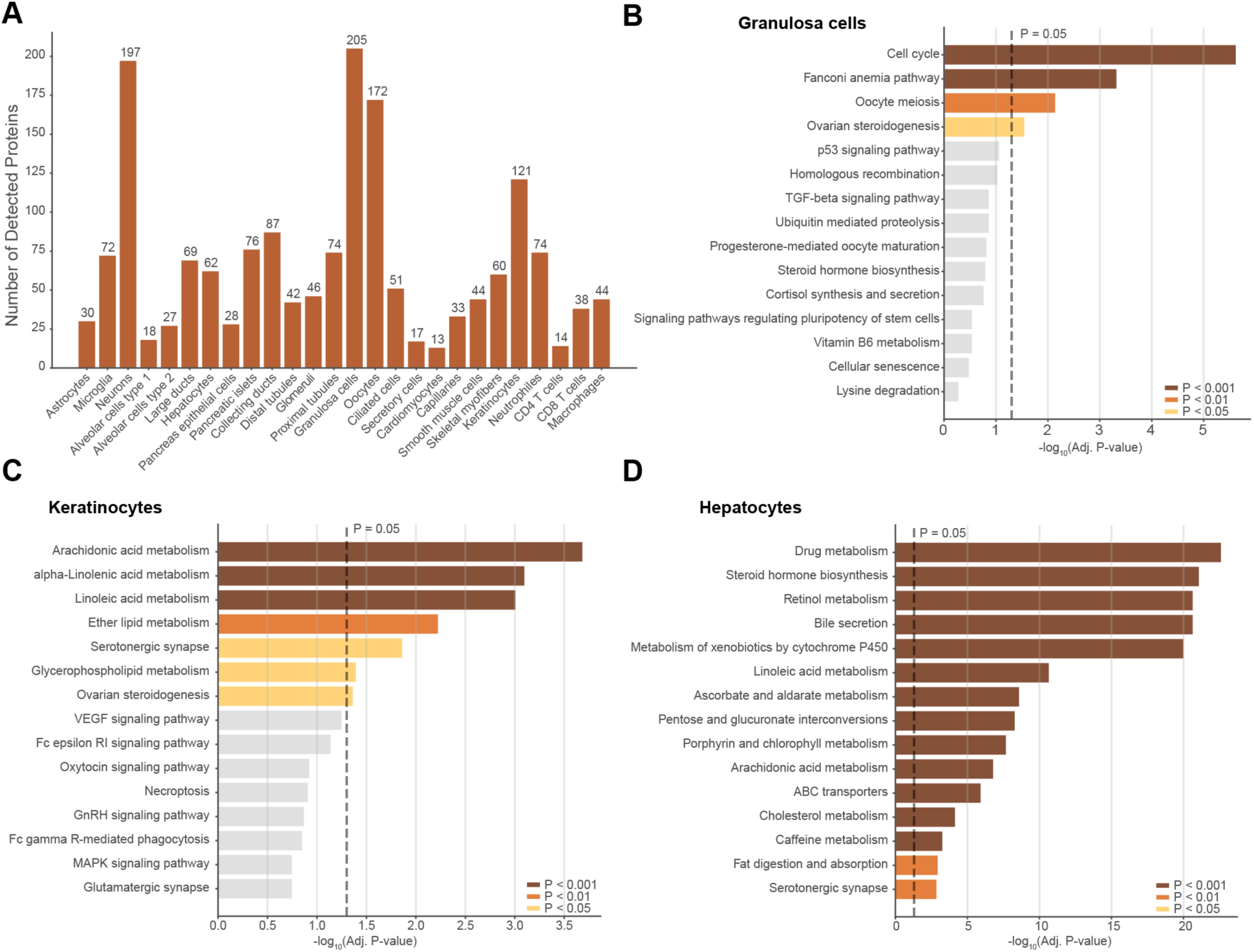
Cell type-specific proteins reveal distinct functional specialization. (**A**) Number of proteins uniquely detected in a single cell type. (**B–D**) KEGG pathway enrichment analysis of cell type-specific proteins for granulosa cells (**B**), keratinocytes (**C**) and hepatocytes (**D**). Bar length indicates - log₁₀(adjusted P-value). Bar color denotes significance level.

**Fig. S11.**
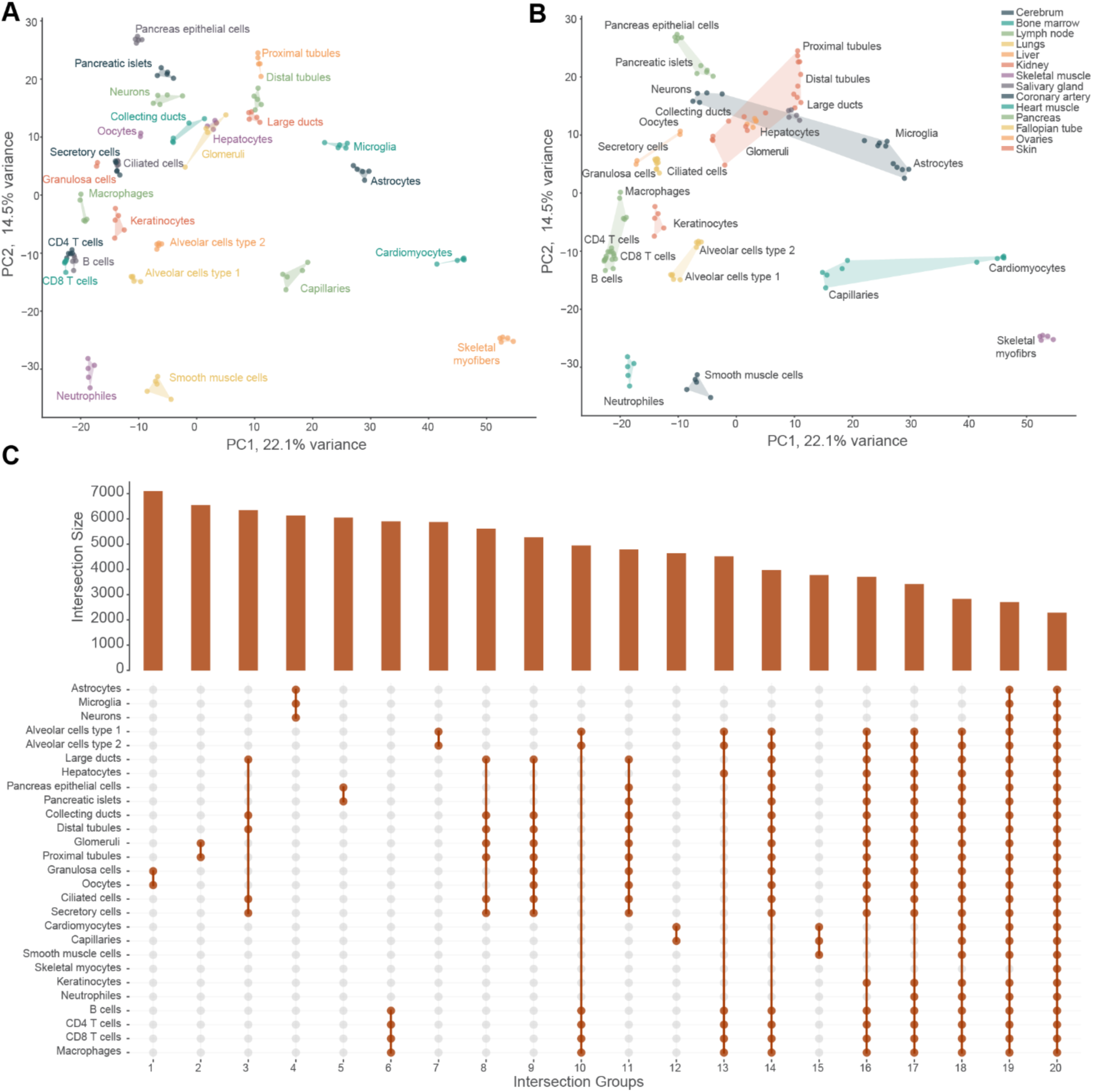
Shared proteome signatures reflect functional relationships across cell types. (**A**) Principal component analysis of the core proteome across all cell types. Individual replicates are color-coded and labeled by cell type. (**B**) Same PCA as in (A) with coloring per tissue of origin. (**C**) UpSet plot showing the top 20 protein intersections across cell types, ranked by intersection size. Bar height indicates the number of shared proteins and connected dots indicate the cell types contributing to each intersection.

**Fig. S12.**
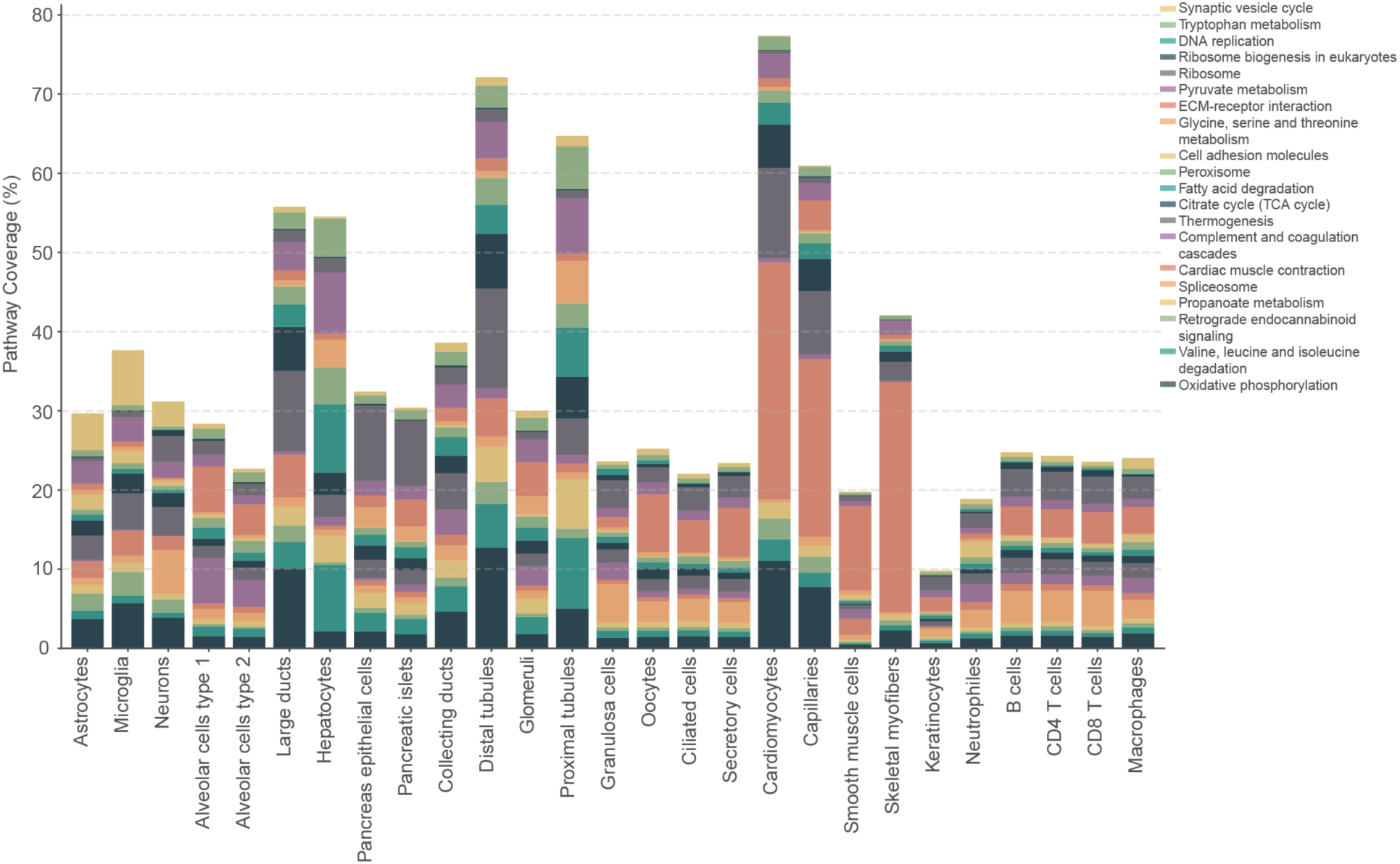
KEGG pathway composition reflects functional specialization across cell types. (**A**) KEGG pathway enrichment analysis across 27 cell types. The 20 pathways with greatest variance across cell types are taken into consideration. Stacked bar plot showing the percentage of total proteome assigned across all cell types.

**Fig. S13.**
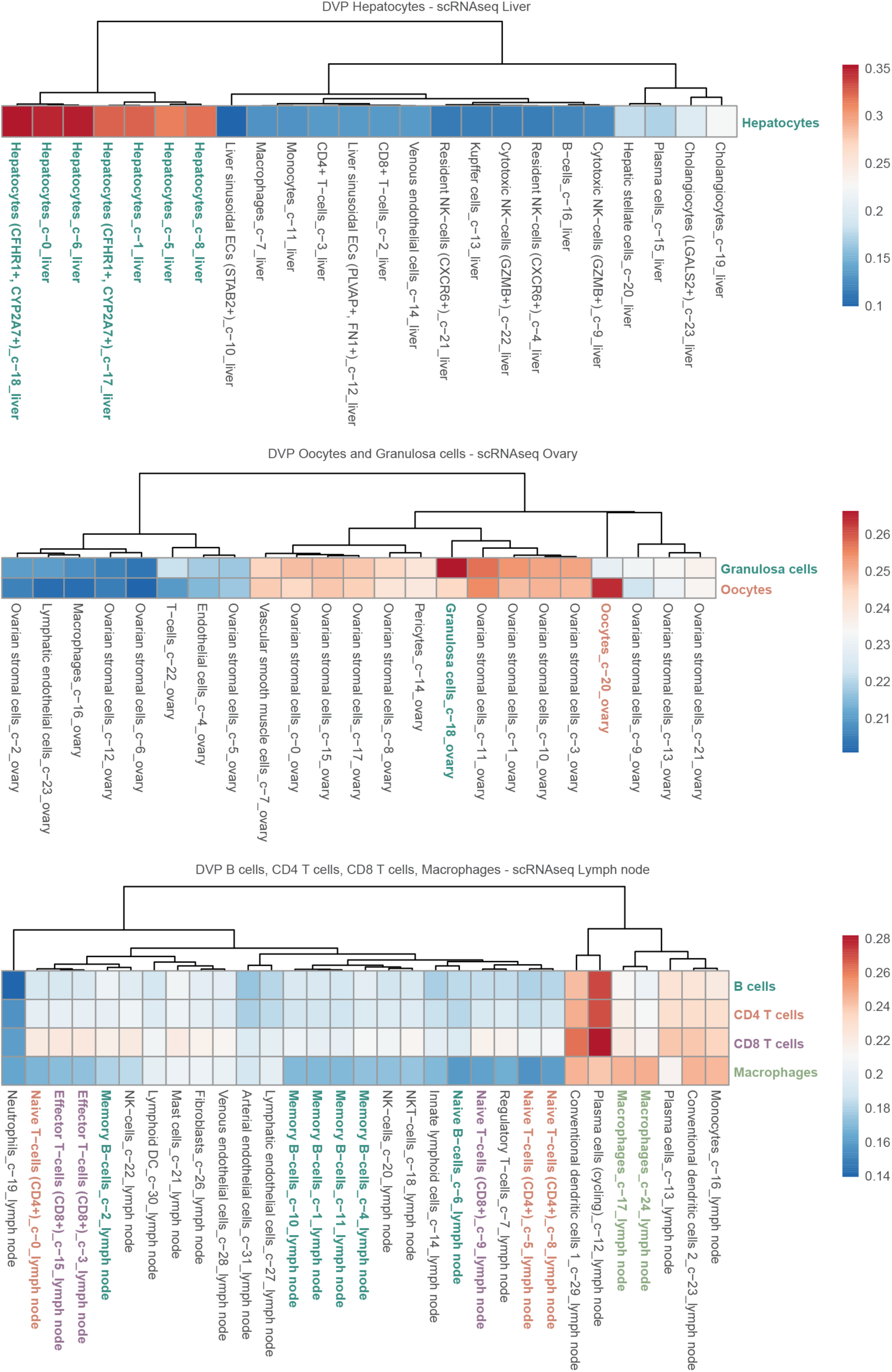
RNA cluster selection for transcriptome-proteome integration. Spearman correlation heatmaps comparing each DVP cell type proteome against all RNA sequencing clusters from the corresponding tissue. Examples shown for hepatocytes (liver), oocytes and granulosa cells (ovary), and B cells, CD4 T cells, CD8 T cells, and macrophages (lymph node).

**Fig. S14.**
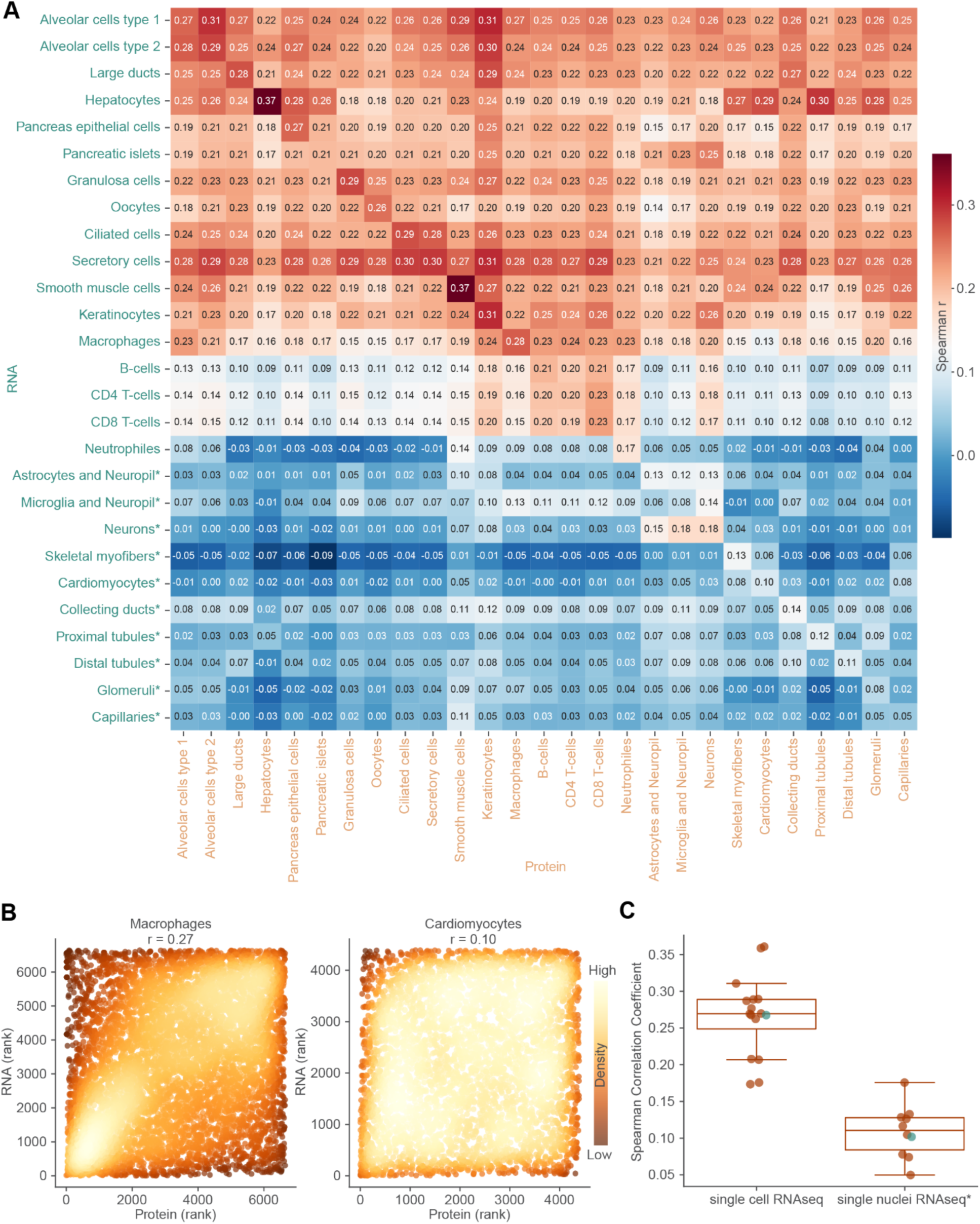
RNA-protein correlation analysis across RNAseq techniques. (**A**) Spearman correlation matrix comparing RNA and protein expression across all cell types (identical to Fig. 4D). Heatmap color and annotations indicate correlation coefficients. Asterisks denote cell types with single-nuclei RNA sequencing data. (**B**) Scatter plots of ranked RNA versus protein expression for matched gene-protein pairs in macrophages (scRNAseq, left) and cardiomyocytes (snRNAseq, right). Spearman correlation coefficients are indicated. (**C**) Comparison of Spearman correlation coefficients between cell types profiled by scRNAseq versus snRNAseq. Green points indicate examples shown in (B). Cell types analyzed by snRNAseq are indicated by an asterisk.

**Fig. S15.**
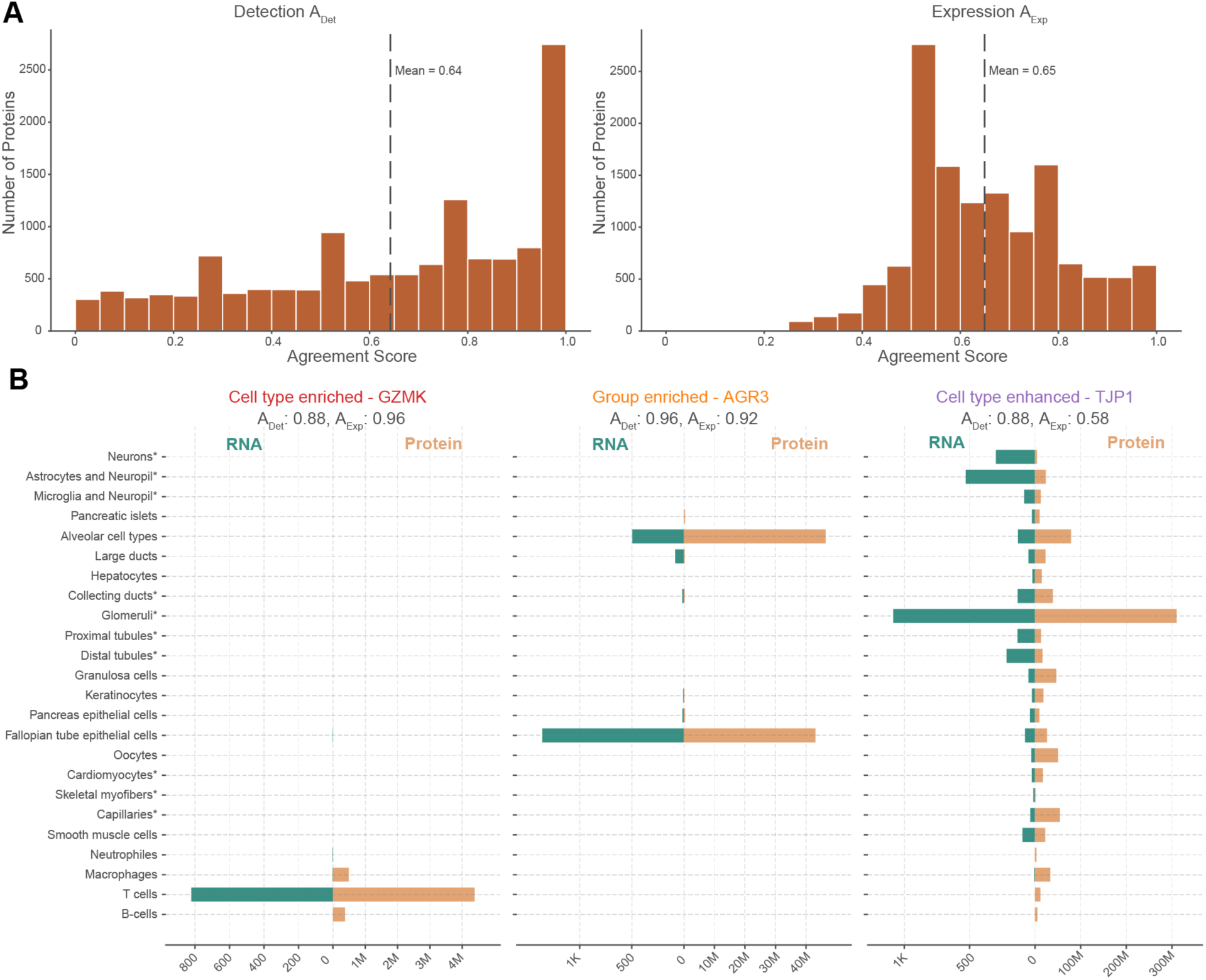
Metrics to evaluate RNA-protein concordance patterns. **(A)** Distribution of detection agreement score (A_Det_, left) and expression agreement score (A_Exp_, right) across all matched gene-protein pairs. Dashed lines indicate mean values. **(B)** Examples illustrating cell type-enriched (GZMK), group-enriched (AGR3), and cell type-enhanced (TJP1) specificity categories. Bar plots show RNA (green, left) and protein (blue, right) expression across all cell types for each example. Cell types analyzed by snRNAseq are indicated by an asterisk

**Fig. S16.**
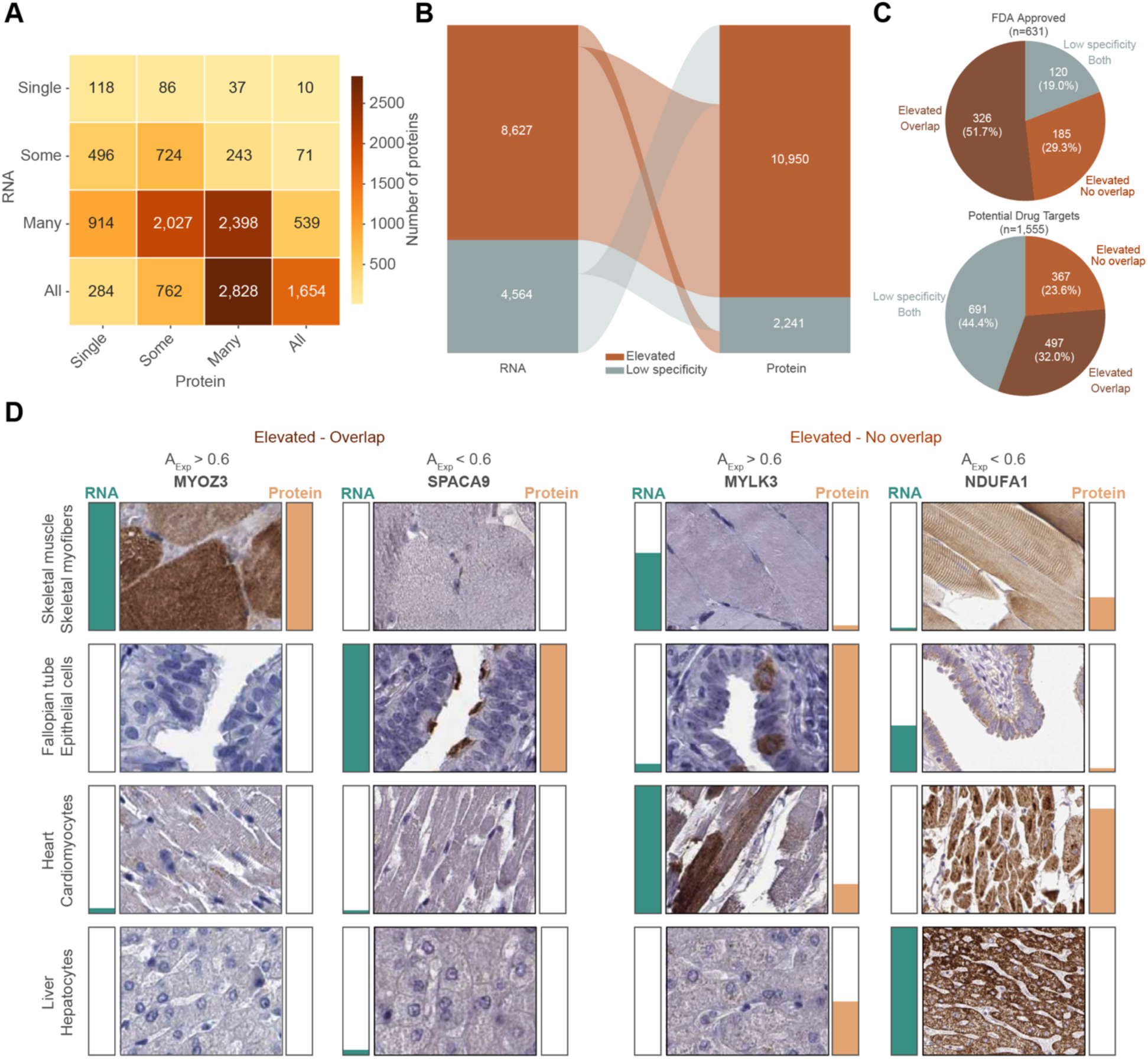
Concordance and divergence of RNA-protein specificity classifications across cell types. **(A)** Heatmap showing the distribution of matched gene-protein pairs across Human Protein Atlas distribution categories in RNA (y-axis) and MS (x-axis). Numbers indicate protein counts per category pair. Color indicates the number of proteins. **(B)** Alluvial diagram showing transitions between elevated (cell type-enriched, group-enriched or cell type-enhanced) and low specificity categories from RNA to protein. Numbers indicate protein counts per category. **(C)** Distribution of elevated overlap, elevated no overlap, and low specificity categories for FDA-approved drug targets (top, n = 635) and potential drug targets (bottom, n = 1,558). Protein classes as annotated by the Human Protein Atlas (see Fig. 2B). **(D)** Representative immunohistochemistry (IHC) images illustrating the four RNA-protein concordance groups defined by elevated overlap status and expression agreement score (A_Exp_). Columns correspond to one representative protein per group: MYOZ3 (elevated overlap, A_Exp_ > 0.6), SPACA9 (elevated overlap, A_Exp_ < 0.6), MYLK3 (elevated no overlap, A_Exp_ > 0.6), and NDUFA1 (elevated no overlap, A_Exp_ < 0.6). Rows show four cell types profiled by DVP: skeletal muscle myofibers, fallopian tube epithelial cells, heart cardiomyocytes, and liver hepatocytes. For each tissue–protein combination, the IHC image is accompanied by bar plots showing normalized RNA expression (teal) and MS protein abundance (orange) across all cell types.

**Fig. S17.**
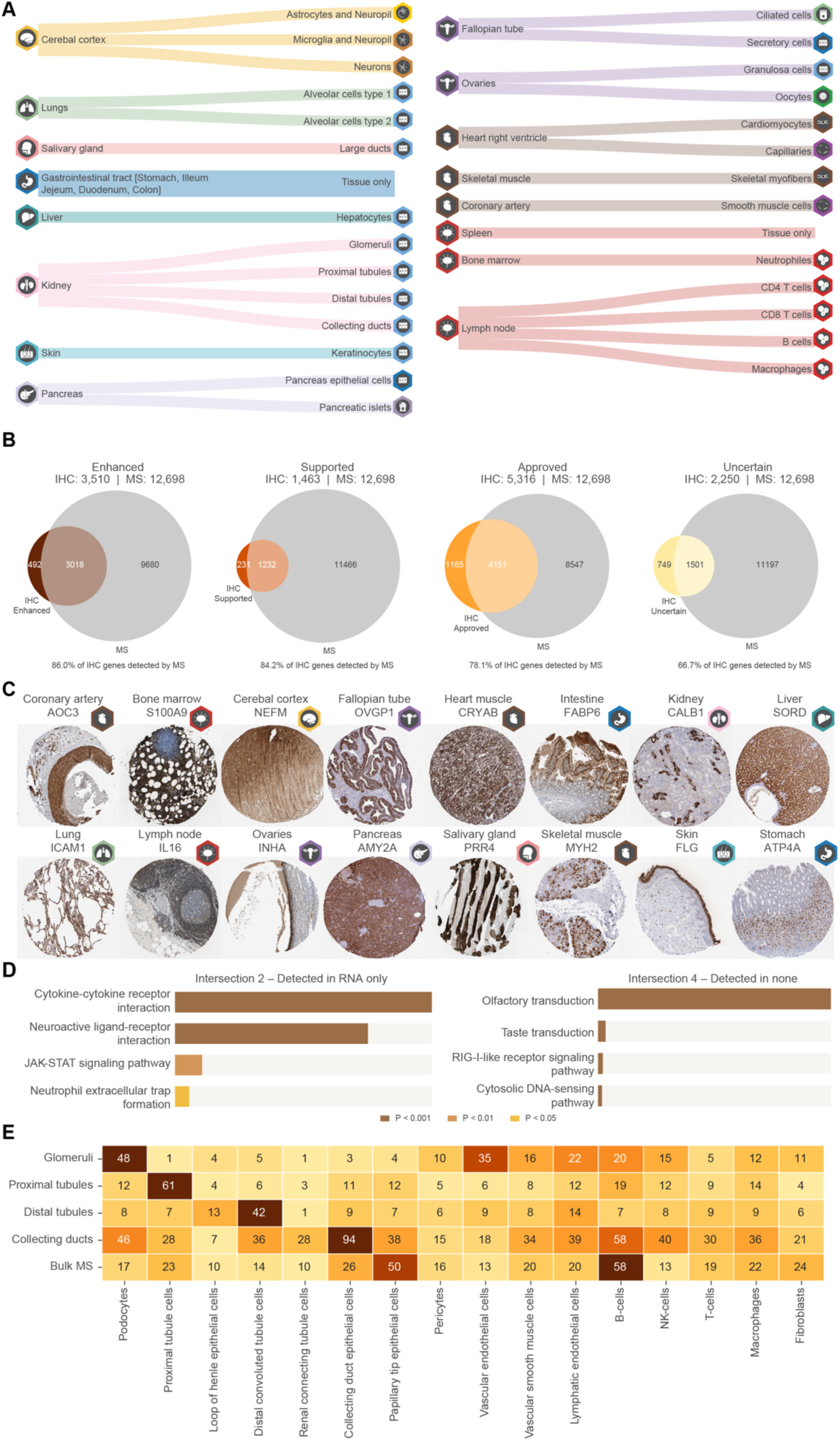
Extension of proteomic atlas coverage through bulk tissue mass spectrometry. **(A)** Overview of the 14 DVP tissues and six additional organs profiled by bulk MS. Icons indicate tissue and cell type category. **(B)** Overlap between HPA IHC-detected proteins and bulk MS detection, shown separately for each reliability tier (Enhanced, Supported, Approved, Uncertain). Circle sizes reflect the total number of proteins per modality; percentages indicate the fraction of IHC-detected proteins recovered by MS. **(C)** Representative IHC images of marker proteins detected concordantly by both IHC and bulk MS across 16 tissues **(D)** Overrepresentation analysis of the top four enriched KEGG pathways for proteins detected exclusively by RNA-based approaches (left) or absent across all modalities (right). Bar color indicates significance level. **(E)** Heatmap showing the number of proteins specific to each kidney DVP cell type or bulk MS, mapped to kidney single-nucleus RNA sequencing clusters. Rows correspond to DVP-profiled kidney cell types and bulk MS; columns correspond to snRNA-seq clusters. Numbers indicate protein counts per MS–cluster pair. Color intensity reflects protein count.

**Fig. S18.**
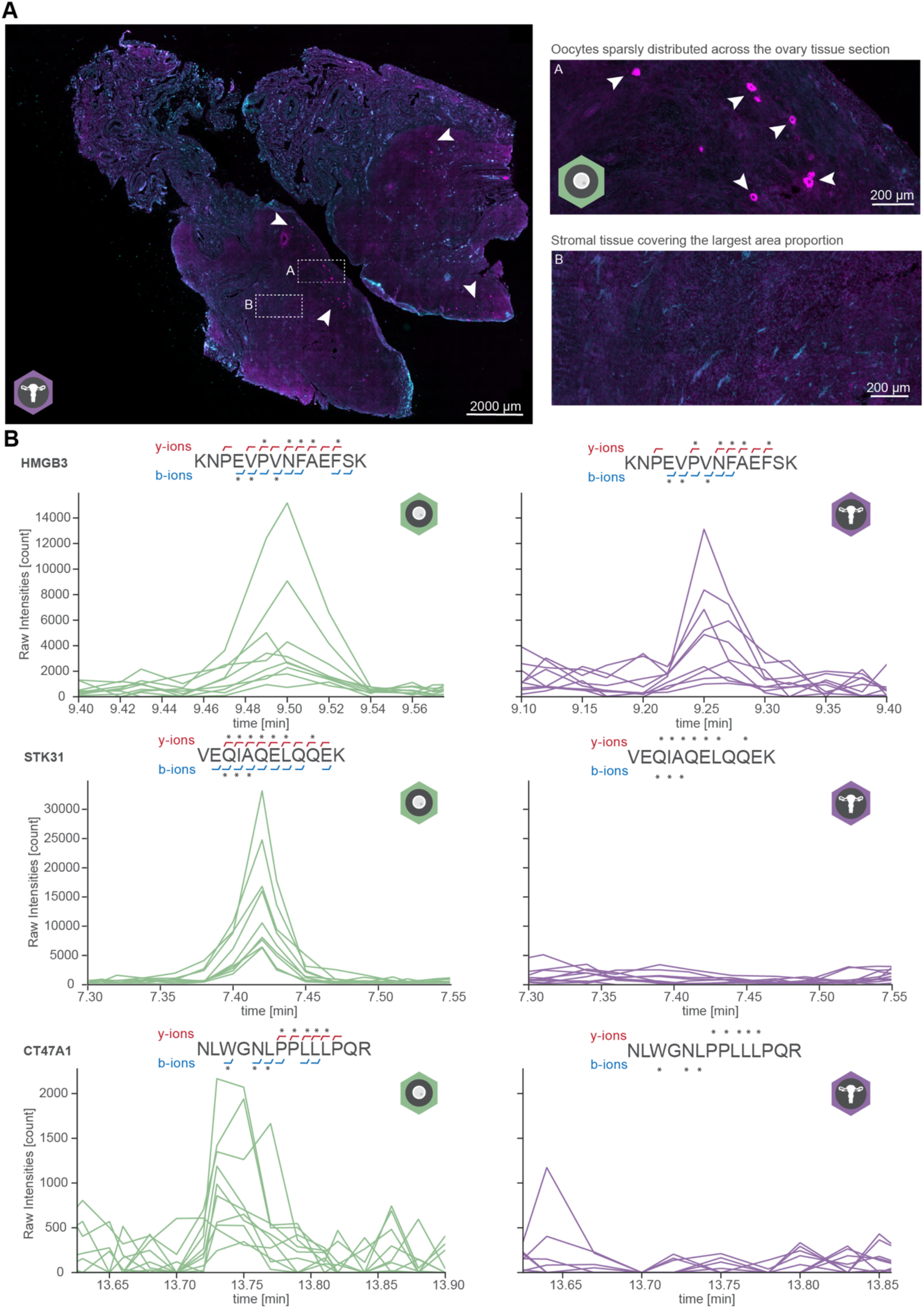
Cell type resolution unmasks cancer-testis antigen detection in oocytes. **(A)** Representative immunofluorescence images of a full ovarian tissue section. Nuclei are shown in cyan, oocyte marker ZP3 in magenta. Arrowheads indicate sparsely distributed oocytes across the tissue section (inset A). Inset B shows the surrounding stromal tissue, which covers the largest area proportion of the section. Scale bars: 2,000 μm (overview), 200 μm (insets). **(B)** Extracted ion chromatograms for representative peptides of three cancer-testis antigens, HMGB3 (KNPEVPVNFAEFSK), STK31 (VEQIAQELQQEK), and CT47A1 (NLWGNLPPLLLPQR), in DVP oocytes (green, left) and bulk ovarian tissue (purple, right). Annotated y-ions (red, above) and b-ions (blue, below) indicate identified fragment ions; asterisks indicate the ion intensity traces plotted in the extracted ion chromatogram. For proteins not detected in bulk, ion annotations are omitted while asterisks are retained to indicate which matching ion traces are displayed. Raw intensity is shown on the y-axis; retention time in minutes on the x-axis.

